# Aggregate-Bound Chaperone Complexes Promote Disaggregation through J-Domain-Driven Hsp70 Recruitment

**DOI:** 10.64898/2026.02.06.704373

**Authors:** Hubert Wyszkowski, Agata Konieczna, Maria Pokornowska, Wiktoria Sztangierska, Filip Kus, Maciej Malolepszy, Katarzyna Kalinowska, Dominik Purzycki, Bartlomiej Tomiczek, Krzysztof Liberek, Agnieszka A. Klosowska

## Abstract

Protein disaggregation by Hsp70 chaperone is essential for maintaining cellular proteostasis, particularly under stress and in proteotoxic diseases. J-domain proteins (JDPs/Hsp40s) interact with Hsp70 through their signature J-domain, stimulating ATP hydrolysis and substrate binding. Disaggregation of stress-related aggregates and amyloid fibrils strictly depends on Class B JDPs. In these specialized co-chaperones, J-domain is autoinhibited and requires activation through JDP interaction with the C-terminal EEVD motif of Hsp70. This mechanism is critical for aggregate disassembly.

Here, we investigated how the two distinctive features of Class B JDPs, EEVD binding and J-domain autoinhibition, promote Hsp70-dependent disaggregation of amorphous aggregates and α-synuclein fibrils. Using purified human and yeast chaperones, we demonstrate that Hsp70 functional cycle with Class B JDPs diverges from the classical model, as EEVD binding stabilizes the JDP-Hsp70-substrate complex. This implies multi-step Hsp70 clustering on aggregates, in which Class B JDPs in complex with aggregate-bound Hsp70 recruit additional Hsp70 molecules to form disaggregation-competent clusters. Experiments with the Hsp70 variant lacking EEVD validate this model but also reveal negative consequences of the stable Hsp70-JDP interaction that make the J-domain autoinhibition indispensable. Collectively, our results highlight EEVD binding and JDP autoinhibition as a coupled regulatory strategy that leads to enhanced Hsp70-mediated protein disaggregation.

## Introduction

Stress induces protein misfolding and aggregation, ultimately leading to proteostasis disruption, a hallmark of ageing and proteotoxic diseases such as Alzheimer’s and Parkinson’s (1). To prevent these effects, cells employ protein quality control systems that rely on molecular chaperones. Among them, the ATP-dependent Hsp70 chaperone plays a key role in counteracting the accumulation of toxic protein aggregates by preventing protein misfolding and by refolding of already misfolded polypeptides (2–4).

Hsp70 chaperone activity relies on co-chaperones: J-domain proteins (JDPs/Hsp40) and nucleotide exchange factors (NEFs)(5, 6). According to the established model, the Hsp70 activity cycle begins when a JDP binds a misfolded protein. Subsequently, the substrate is transferred to Hsp70 and the J-domain of the JDP binds at the interface between the Nucleotide Binding (NBD) and Substrate Binding (SBD) domains of Hsp70. JDP and substrate binding stimulates ATP hydrolysis, which induces conformational changes in Hsp70, resulting in tight closure of the SBD on the misfolded protein. These rearrangements disrupt the NBD/SBD interface, leading to J-domain dissociation from the ADP-bound Hsp70 (2, 7–10). The substrate remains bound to Hsp70 (11) and can be released only upon ADP to ATP exchange, a slow process that is accelerated by NEFs, such as Hsp110 (12–14). Once released, the polypeptide substrate may successfully fold or enter another round of chaperone-assisted processing.

Hsp70-dependent protein folding and disaggregation in the cytosol and nucleus are facilitated by evolutionarily conserved JDPs assigned to Class A and Class B, which share common structural elements: the N-terminal J-domain, followed by a mostly unstructured glycine/phenylalanine-rich region (G/F), two beta-sandwich CTD domains involved in substrate binding (CTD1 and CTD2) and a C-terminal dimerization domain. Class A features an additional Zinc Finger-Like Motif (ZnF) in its CTD1. Despite their considerable structural similarities, Class A and Class B JDPs demonstrate clearly divergent activities (15–19).

Class A JDP proteins specialize in early recognition and capture of misfolded polypeptides, preventing aggregation and promoting their folding and delivery to Hsp70 (20, 21). In contrast, Class B proteins possess a unique ability to facilitate solubilization of amorphous and amyloid aggregates by organizing packed Hsp70 assemblies at the aggregate surface (22–25). Such Hsp70 clusters favor aggregate dissolution through entropic pulling; a mechanism whereby trapped polypeptides are extracted from aggregates by the thermodynamic drive toward greater conformational freedom (26–28). Class B JDP-dependent disaggregation activity of the Hsp70 system is required not only for protein recovery, but also for aggregate fragmentation that precedes downstream processing by proteasome and autophagy (29).

Recent studies highlight how several evolutionary changes in JDPs have potentiated Hsp70 as a disaggregating system. These adaptations in Class B ancestors include the loss of the ZnF region, G/F elongation, autoinhibition of the J-domain by Helix 5 (H5) localized within the G/F and the auxiliary interaction with the EEVD motif of Hsp70 (16). As a result, an effective Class B JDP-Hsp70 interaction involves two binding events. The first binding occurs between the CTDI domain and the EEVD motif of at the C-terminus of Hsp70’s SBD (30–32). This interaction leads to H5 dissociation from the J-domain and the release of autoinhibition (23, 33–35). This enables the second, canonical binding between the J-domain and the NBD/SBD interface of Hsp70, which triggers ATP hydrolysis.

Hsp70’s disaggregation activity strictly depends on Class B JDP autoinhibition by Helix 5 (23, 33, 35). Disruption of this inhibition by mutations within the J-domain or G/F impairs Hsp70 clustering on aggregate surfaces and compromises disassembly of α-synuclein fibrils (22, 23). Defective autoinhibition has also been linked to myopathies. Among clinically relevant mutations identified in the Class B JDP paralog DNAJB4 and a non-canonical Class B member DNAJB6, some affect the J-domain– H5 interface and lead to the accumulation of protein aggregates and muscle pathology, possibly through Hsp70 dysregulation (23, 36–38).

Consequently, deletion of the EEVD motif completely abolishes Hsp70 activity when paired with Class B JDPs (22, 23, 30, 31, 33, 35). Engagement of the Hsp70 EEVD motif relieves Class B JDP from its autoinhibited state. Mutations that unlock the J-domain partially restore protein folding by Hsp70 variants lacking EEVD (ΔEEVD) (23, 33, 35) but fail to recover their disaggregation activity. Therefore, Class B JDP variants with disrupted autoinhibition in combination with Hsp70^ΔEEVD^ resemble Class A JDPs: supporting refolding of non-aggregated substrates but ineffective in disaggregation of amorphous aggregates (22) and amyloid fibrils (23). The structural basis of Class B JDP activation upon EEVD binding and its role in protein disaggregation remain unclear.

Another consequence of EEVD binding is the stabilization of the Hsp70-JDP complex. Our recent findings demonstrate that the interaction between the yeast Hsp70 Ssa1 and Class B JDP Sis1 via EEVD remains stable even when Hsp70 is in the ADP state (22). This suggests that the model of the Hsp70 cycle, largely based on bacterial DnaJ protein categorized as a Class A JDP, does not represent Class B JDPs, thereby obscuring our understanding of JDP mechanisms in Hsp70-dependent disaggregation.

Here, we investigate how the distinctive features of Class B JDPs, Hsp70 interaction mediated by the EEVD motif and autoinhibition of the J-domain, synergize to drive the formation of chaperone clusters at the surface of aggregated proteins. We show that Class B JDP interaction with EEVD stabilizes the JDP-Hsp70-aggregate complex. We explore the potential role of such complexes in cluster formation and unravel the secondary, EEVD-independent mode of Hsp70 recruitment to aggregates, which potentiates the disaggregation activity of the Hsp70 system. On the other hand, we show that JDP binding through EEVD may limit the fraction of Hsp70 molecules in the ATP-bound state competent for substrate engagement, thereby imposing the dependence of efficient disaggregation on J-domain autoinhibition. Together, our results reveal interdependent mechanisms that enable Class B JDPs to promote effective disaggregation.

## Results

### Class B JDP interaction with EEVD reshapes Hsp70 activity cycle

Class B JDP binding to Hsp70 is a two-step process. In addition to the J-domain-mediated interaction, which is disrupted when Hsp70 hydrolyses ATP to ADP, Class B JDPs interact with the C-terminal EEVD motif of Hsp70 (Fig. 1A). We sought to determine whether this additional interaction is maintained after ATP hydrolysis, when Hsp70 is in the ADP-bound state and in complex with a substrate.

**Figure 1.**
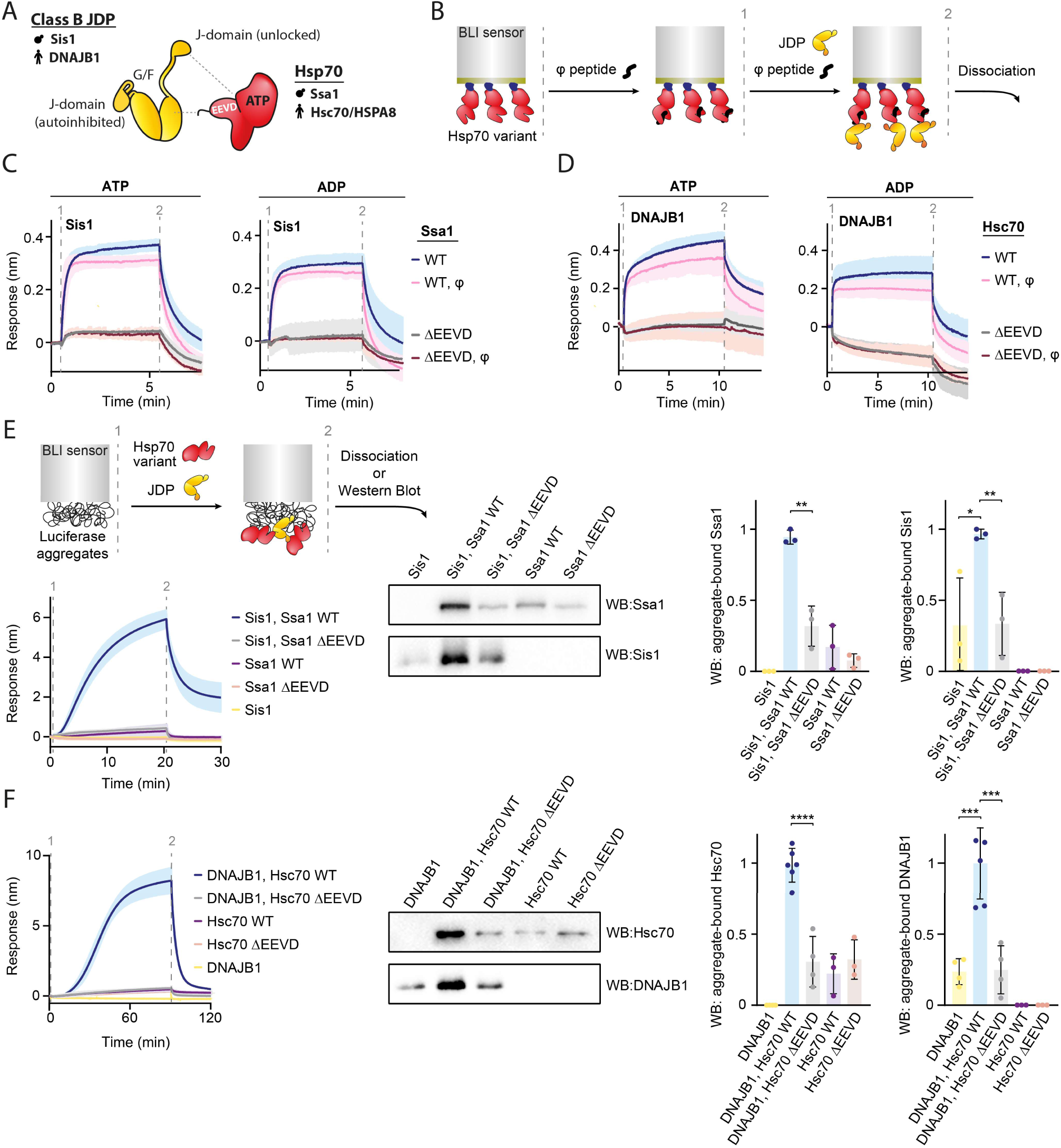
Interaction with EEVD stabilizes JDP complex with substrate-bound Hsp70. (A) Class B JDP interaction with Hsp70 involves two binding sites (dashed lines). JDP binding to the EEVD motif of Hsp70 releases the J-domain from autoinhibition, allowing the J-domain to bind to the ATP-bound Hsp70. (B) Experimental scheme for panels C, D. Yeast or human Hsp70 variant was immobilized on the BLI sensor through the His-SUMO tag and incubated for 30 min with or without the ϕ peptide (FYQLALT) in the presence of ATP (1 mM) or ADP (1 mM), following the addition of Class B JDP to the reaction mixture and further incubation for 5 min. Dissociation was initiated by transferring the sensor to the same buffer without JDPs. (C) Interaction of Ssa1 (WT or ΔEEVD) with Sis1 (0.4 µM) in the presence of ATP (left panel) or ADP (right panel), with or without the ϕ peptide (10 µM). (D) Hsc70 interaction with DNAJB1 (2 µM) in the presence of ATP (left panel) or ADP (right panel), with and without the ϕ peptide (10 µM). (E) Upper panel: experimental scheme for panels E and F. BLI sensor with luciferase aggregates was incubated with yeast or human Hsp70 variants and Class B JDPs in the presence of ATP (5 mM). Upon binding, sensors were either incubated with buffer lacking chaperones to follow protein dissociation or subjected to Western Blot analysis (WB). Lower panels: EEVD motif in Ssa1 facilitates Sis1 association with aggregates. Left panel: Binding of Ssa1 WT or ΔEEVD (1 µM) and Sis1 (1 µM) to sensor-bound luciferase aggregates. Middle panel: Western Blot analysis of Ssa1 (upper blot) and Sis1 (lower blot) bound to the sensor after 20 min incubation with the indicated chaperones. Right panels: Normalized Western Blot signals: the level of Ssa1 and Sis1 bound to luciferase aggregates. (F) The EEVD motif in Hsc70 is critical for DNAJB1 association with aggregates. Left panel: Binding of Hsc70 WT or ΔEEVD (2 µM) with DNAJB1 (2 µM) to sensor-bound luciferase aggregates. Middle panel: Western Blot analysis of Hsc70 (upper blot) or DNAJB1 (lower blot) bound to the sensor after 1.5 h of incubation with the indicated chaperones. Right panels: Normalized Western Blot signals: the level of Hsc70 and DNAJB1 bound to luciferase aggregates. (C-F) Binding curves represent the mean of at least three replicates, with shades indicating SD. Dashed line 1 indicates the start of the binding step, dashed line 2 indicates the dissociation step into buffer lacking chaperones. (E, F) The presented blots are representative for at least 3 repeats. Signal intensities were normalized to one of the Hsp70^WT^-JDP replicates. Error bars represent SD. Unpaired T test: *p<0.05, **p<0.01, ***p<0.001, ****p<0.0001

We chose the yeast Hsp70-JDP system Ssa1-Sis1, as Sis1 is the sole Class B JDP in *S. cerevisiae* (16) and is therefore likely to represent the general properties of this JDP Class without the confounding effects of paralog-specific functional specialization. In parallel, we examined the human pair Hsc70(HSPA8)-DNAJB1, as these paralogs are the most effective in disaggregation, system with robust activity toward amyloid fibrils (24, 25, 39–41).

To monitor Hsp70-Class B JDP interaction, we employed Bio-Layer Interferometry (BLI), which allows real-time measurements of protein layer thickness on the surface of an optical sensor. We immobilized Hsp70 (Ssa1 or Hsc70) on the BLI sensor via an N-terminal His-SUMO tag (Supplementary Fig. 1A). Subsequently, we incubated the sensor-bound Hsp70 with or without an excess of the model Hsp70 substrate, ϕ peptide (FYQLALT) (42, 43) for 30 min, to saturate the substrate-binding domain of Hsp70. Such a small substrate was chosen to prevent simultaneous binding of both Hsp70 and the JDP to the same substrate molecule.

Upon subsequent incubation of sensor-bound Hsp70 with the corresponding JDP (Sis1 or DNAJB1), the BLI signal increased, indicating formation of an Hsp70-JDP complex (Fig. 1B-D). JDP binding to Hsp70 was comparable in the presence of either ATP or ADP throughout the experiment and was only slightly reduced in the presence of ϕ peptide (Fig. 1C,D). This suggests that Class B JDPs efficiently bind to ADP-bound, substrate-engaged Hsp70.

Hsp70 variants lacking the EEVD motif, Ssa1^ΔEEVD^ and Hsc70^ΔEEVD^, exhibited only marginal JDP binding (Fig. 1C,D). This is in agreement with earlier findings that the Hsp70-JDP interaction mediated only through the J-domain is hardly detectable by conventional protein-protein interaction assays (10, 22). The observed EEVD-dependent JDP binding is consistent with the established key role of the EEVD motif in Hsp70 collaboration with Class B JDPs (19, 30, 31, 33, 35).

Hsp70 variant deficient in ATP hydrolysis, Ssa1^T201A^, exhibited JDP-binding signal similar to that of the wild-type Ssa1 (Ssa1^WT^) (Supplementary Fig. 1B), further supporting the hypothesis that the Hsp70-Class B JDP interaction via EEVD is uncoupled from the ATPase cycle of Hsp70. The dependence of the Ssa1^T201A^-Sis1 interaction on the EEVD motif was supported by both BLI and MST analyses (Supplementary Fig. 1B,C,D).

We next explored functional consequences of the ability of Class B JDPs to interact with the EEVD motif of substrate-bound Hsp70. During protein disaggregation, Hsp70 assembles into clusters on the aggregate surface in Class B JDP-dependent manner (22–25). We asked whether Hsp70, in turn, promotes Class B JDP recruitment to the aggregates via interaction with EEVD.

To address that, we analyzed aggregate binding by yeast and human orthologs: wild-type Hsp70 (Ssa1^WT^ or Hsc70^WT^) and their variants lacking EEVD (Ssa1^ΔEEVD^ or Hsc70^ΔEEVD^), in combination with JDPs (Sis1 and DNAJB1). We prepared BLI-sensors with heat-aggregated luciferase (Supplementary Fig. 2). The sensors were subsequently incubated with the chaperone mixtures. Based on the increase in the BLI signal, both the yeast and human Hsp70-JDP systems bound efficiently to luciferase aggregates, provided that both Hsp70 and JDP were present together, and that the EEVD motif of Hsp70 was intact (Fig. 1E,F). Western Blot analysis of the individual chaperones associated with the aggregates on the BLI sensors showed that Hsp70 promoted JDP association with aggregates in an EEVD-dependent manner (Fig. 1E,F).

Overall, these results show that the interaction of Class B JDPs with the EEVD motif of Hsp70 stabilizes the JDP-Hsp70-substrate complex and facilitates the mutually dependent association of Hsp70 and JDP with aggregates. This suggests that chaperone assemblies formed on the aggregate surface comprise not only Hsp70, but also Class B JDP molecules associated with Hsp70 through the EEVD motif.

### Multi-step Hsp70 binding to protein aggregates

Given that Class B JDP molecules are incorporated into the Hsp70-aggregate complex through the interaction with the EEVD motif, their J-domains are expected to be released from autoinhibition (23). This may allow the activated J-domains within the initial complex on the aggregate to recruit additional Hsp70 molecules, thereby promoting Hsp70 clustering on aggregate surface.

To explore such multi-step Hsp70 binding mechanism, we used the Ssa1^ΔEEVD^ variant, which cannot relieve J-domain autoinhibition and is therefore impaired in aggregate binding and disaggregation with Class B JDPs (Figs. 1E, 2 A-D)(22). We examined whether Ssa1^ΔEEVD^ could nevertheless be recruited to aggregates through the J-domains of Sis1 activated in the presence of wild-type Ssa1. When aggregate-covered BLI sensors were simultaneously incubated with Sis1 and a mixture of Ssa1^WT^ and Ssa1^ΔEEVD^ variants, the binding was more rapid and reached higher levels than with the Ssa1^WT^-Sis1 pair (Fig. 2A). We observed similar effects with another substrate, comprising yeast cell lysate proteins heat-aggregated on the BLI sensor. In this case, the presence of Ssa1^ΔEEVD^ also stimulated the Ssa1^WT^-Sis1 system (Supplementary Fig. 3A), suggesting a common mechanism of Hsp70 binding to different aggregated substrates.

**Figure 2.**
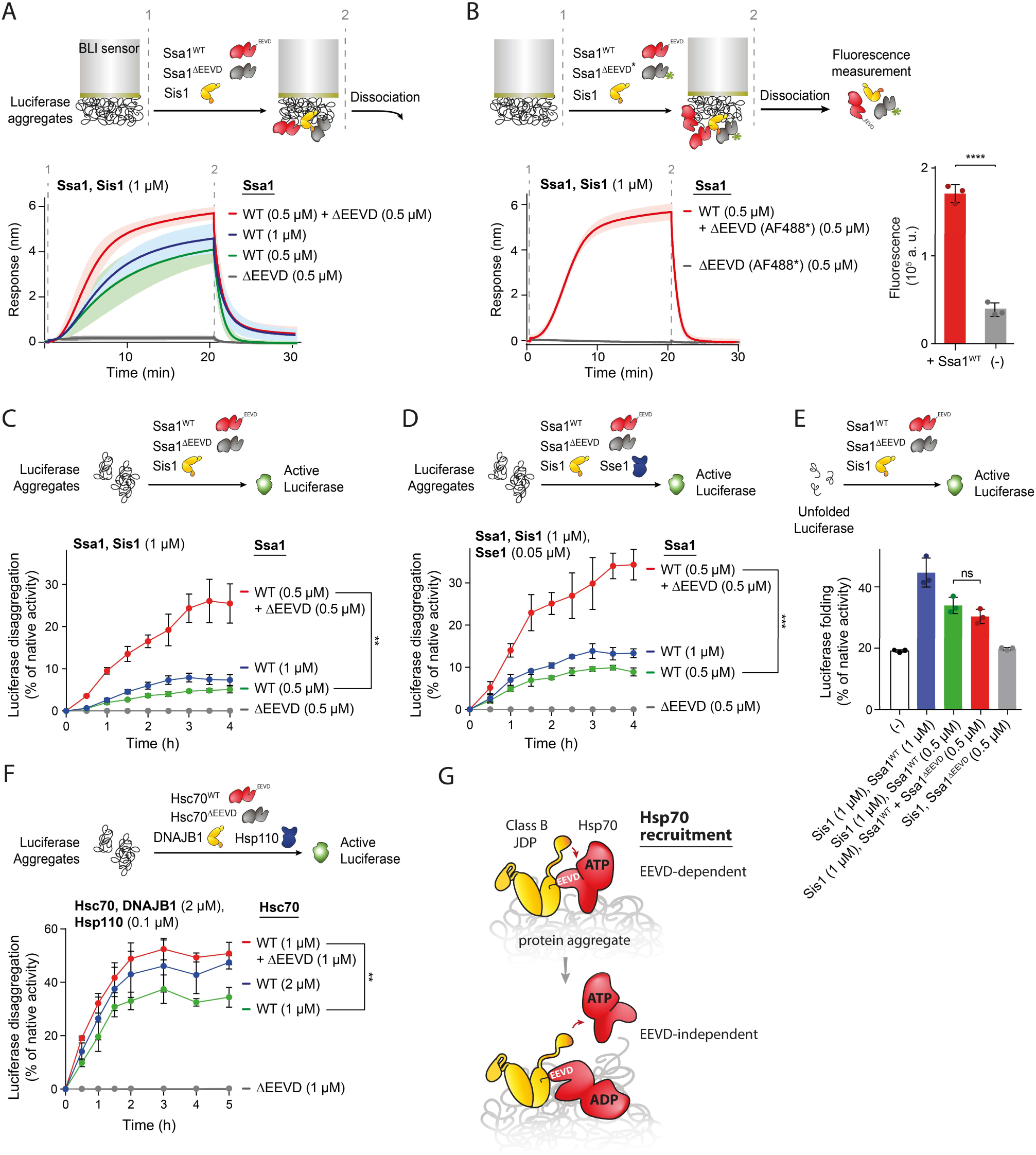
EEVD-dependent and independent collaboration between Hsp70 and Class B JDP in aggregate binding and disaggregation. (A) Ssa1^ΔEEVD^ stimulates chaperone binding to the aggregate when wild-type Ssa1 and Sis1 are present. BLI sensor with aggregated luciferase was incubated with Sis1 and the indicated Ssa1 variants: Ssa1^WT^ (green and blue), 1:1 mixture of Ssa1^WT^ and Ssa1^ΔEEVD^ (red) or Ssa1^ΔEEVD^ alone (grey). (B) Ssa1^ΔEEVD^ binds to aggregates in the presence of Ssa1^WT^ and Sis1. BLI sensor with aggregated luciferase was incubated with Sis1 and AF488-labeled Ssa1^ΔEEVD^ alone (grey) or additionally with Ssa1^WT^ (red). Right panel: Fluorescence intensity of Ssa1^ΔEEVD*AF488^ dissociated from BLI sensor with luciferase aggregates. Unpaired T test: ****p<0.0001. (A, B) Binding curves represent the mean of three experiments, with shaded areas indicating SD. Dashed line 1 indicates the start of the binding step, dashed line 2 indicates the dissociation step into buffer lacking chaperones. (C, D) Ssa1^ΔEEVD^ stimulates disaggregation of luciferase aggregates in the presence of Ssa1^WT^-Sis1 (C) and Ssa1^WT^-Sis1-Sse1 (D). Luciferase aggregates were incubated with the indicated chaperone mixtures and luciferase activity was measured at indicated time points and normalized to native protein. (E) Ssa1^ΔEEVD^ does not stimulate folding of unfolded, non-aggregated luciferase in the presence of Ssa1^WT^-Sis1. GuHCl-unfolded luciferase was diluted from the denaturant directly into the chaperone mixtures. The activity was measured after 1 h of incubation with the indicated chaperone mixtures or without chaperones. (F) Disaggregation of aggregated luciferase by DNAJB1, Hsp105 and the indicated Hsp70 variants: Hsc70^WT^ (blue and green), 1:1 mixture of Hsc70^WT^ and Hsc70^ΔEEVD^ (red) or Hsc70^ΔEEVD^ (grey). (A-F) Upper panels: experimental scheme. (C-F) Error bars indicate the mean ± SD from at least three independent experiments. Unpaired T test was performed for final luciferase activity **p<0.01, ***p<0.001, ns non-significant (G) Proposed multi-step mechanism of Hsp70 binding to aggregates. Upon formation of the initial Class B JDP-Hsp70-aggregate complex, in which the JDP interacts with the EEVD motif of aggregate-bound Hsp70, the unlocked J-domain may recruit additional Hsp70 molecules independently of their EEVD motifs.

Interestingly, when sensor-immobilized luciferase aggregates were initially incubated with Ssa1^WT^ and Sis1, subsequent addition of Ssa1^ΔEEVD^ to the chaperone mixture resulted in substantially higher binding levels than addition of an equivalent amount of Ssa1^WT^ (Supplementary Fig. 3B). However, the effect of Ssa1^ΔEEVD^ was observed only when Ssa1^WT^ was maintained throughout the experiment. When the chaperone mixture present in the second step was lacking Ssa1^WT^, the chaperones rapidly dissociated from the aggregate surface (Supplementary Fig. 3B). This suggests that EEVD-independent Hsp70 recruitment to aggregates by Class B JDPs can be efficient only through continued cooperation with wild-type Hsp70.

To verify whether Hsp70 lacking EEVD is indeed incorporated into the chaperone complexes on the aggregate, we fluorescently labelled Ssa1^ΔEEVD^ with Alexa Fluor 488 (Ssa1^ΔEEVD*AF488^) and incubated it with Sis1 and sensor-immobilized luciferase aggregates (Fig. 2B). We measured the fluorescence of proteins initially bound to aggregates following dissociation step. The mixture of Ssa1^WT^ and Ssa1^ΔEEVD*AF488^ variants yielded five times stronger fluorescence compared to Ssa1^ΔEEVD*AF488^ without Ssa1^WT^ (Fig. 2B), suggesting that Ssa1^ΔEEVD*AF488^ was bound to the aggregates due to the presence of Ssa1^WT^. We next assessed whether the two Ssa1 variants, Ssa1^WT^ and Ssa1^ΔEEVD^, form clusters on protein aggregates. We used a FRET assay, with each Hsp70 variant fluorescently labelled (Donor: Ssa1^ΔEEVD*AF488^; Acceptor: Ssa1^WT*AF594^), incubated together with luciferase aggregates. An increase in the FRET efficiency upon the addition of Sis1 (Supplementary Fig. 3C) suggests that the two Ssa1 variants, wild-type and ΔEEVD, are in close proximity upon JDP-dependent aggregate binding. Together, these findings show that Hsp70 variants lacking the EEVD motif can be recruited to aggregates through cooperation with EEVD-containing Hsp70 ^WT^ molecules and Class B JDPs.

To determine whether, and to what extent, the increased Hsp70 binding to aggregates observed in the presence of the Hsp70^ΔEEVD^ variant (Fig. 2A, Supplementary Fig. 3A,B) correlates with the recovery of active protein, we measured disaggregation of aggregated luciferase by Ssa1^WT^ and Sis1, either alone or in combination with the Ssa1^ΔEEVD^ variant. The presence of Ssa1^ΔEEVD^ stimulated the disaggregation by Ssa1^WT^-Sis1 (Fig. 2C). The effect was even more pronounced at higher concentrations of the chaperones (Supplementary Fig. 3D). The stimulatory effect of Ssa1^ΔEEVD^ on Ssa1 ^WT^-Sis1 activity was also observed in the presence of Hsp110 nucleotide exchange factor Sse1 (Fig. 2D). Notably, with Class A JDP Ydj1, enhanced disaggregation activity with Ssa1^ΔEEVD^ versus Ssa1^WT^ was not observed (Supplementary Fig. 2E). Together, these results suggest that the recruitment of additional Hsp70 molecules via the EEVD-independent, Class B JDP-specific mechanism improves protein disaggregation.

We next asked whether the EEVD-containing Hsp70 molecule needs to be able to bind a substrate to unlock the collaboration between Class B JDP and the Hsp70^ΔEEVD^ variant, or the activation of the J-domain by the EEVD motif may occur already in the solution. To assess that, we used the Ssa1^V435F^ mutant defective in substrate binding (44), as the sole source of Ssa1 containing the EEVD motif. In combination with Sis1^WT^, the Ssa1^V435F^ variant was unable to bind aggregates or recover aggregated luciferase, regardless of the Ssa1^ΔEEVD^ presence (Supplementary Fig. 4A,B). In a similar experiment, we used EEVD-containing Ssa1 variant T201A deficient in ATP hydrolysis, which forms only transient complexes with protein substrates (45). Ssa1^T201A^ promoted very low level of Sis1-Ssa1^ΔEEVD^ binding to the aggregate on the BLI sensor and did not support the disaggregation activity (Supplementary Fig. 4A,B). This suggests that Hsp70 molecules can be recruited to disaggregation independently of their EEVD motifs only by the aggregate-associated Hsp70^WT^-Class B JDP complex.

The EEVD-independent Hsp70 binding by Hsp70^WT^-Class B JDP was effective in the case of aggregates, which present large surfaces rich in chaperone-binding sites. To verify, whether Ssa1^ΔEEVD^ also improves the recovery of smaller and soluble misfolded protein substrates, we analyzed the protein folding activity of the Hsp70-JDP system with non-aggregated, unfolded luciferase. GuHCl-unfolded luciferase was extensively diluted from a denaturant directly into the chaperone mixture to avoid aggregate formation. In contrast to the disaggregation, no stimulation of the protein folding activity of Ssa1^WT^-Sis1 by the Ssa1^ΔEEVD^ variant was observed (Fig. 2E).

Finally, to assess whether the potentiation of the Hsp70 system by the ΔEEVD variant is conserved, we examined the effect of human Hsc70^ΔEEVD^ variant on the disaggregation activity of Hsc70^WT^ in the presence of DNAJB1. We performed luciferase disaggregation according to the protocol previously established for the human system (12), including the Hsp110 NEF (Hsp105) in the reaction due to the stronger dependence of the human disaggregation system on nucleotide exchange factors (12, 46). Similarly to the yeast system, the mixture of Hsc70^WT^ and Hsc70^ΔEEVD^ in the presence of DNAJB1 and Hsp110 exhibited higher disaggregation activity than the reaction with Hsc70^WT^ as theonly Hsp70 variant (Fig. 2F). However, the stimulation by the ΔEEVD variant was less pronounced than that observed for the yeast system.

The above results suggest that Hsp70 collaborates with Class B JDPs in aggregate binding and disaggregation through two interdependent mechanisms: one dependent on, and one independent of, the interaction between the JDP and the EEVD motif of Hsp70 (Fig. 2G). The J-domain activated within the initial EEVD-dependent Hsp70-JDP complex that forms on aggregates appears to be sufficient to recruit additional Hsp70 molecules in close proximity, without involvement of their EEVD motifs.

### Class B JDP-Hsp70 interaction via EEVD limits Hsp70 activity

It appears paradoxical that when half of Hsp70 in the mixture with Class B JDP is replaced by the otherwise inactive Hsp70^ΔEEVD^ variant, aggregate binding and disaggregation activity are enhanced (Fig. 2A,C,D,F; Supplementary Fig. 3A,B,D). This suggests that presence of the EEVD motif prevents the wild-type Hsp70 paired with Class B JDP from reaching its full potential. To investigate this, we analyzed the effects of Hsp70^ΔEEVD^ across a range of Class B JDP concentrations. At low Sis1 concentration (0.1 µM), the Ssa1^WT^ and Ssa1^ΔEEVD^ mixture exhibited much slower aggregate binding and less efficient disaggregation than Ssa1^WT^ only (Fig. 3A,B, Supplementary Fig. 5A,B). This suggests that at low JDP levels, an EEVD-dependent step becomes rate-limiting. In contrast, higher Sis1 concentrations (>0.5 µM) favored the stimulation by Ssa1^ΔEEVD^ and inhibited the system with Ssa1^WT^ as the sole Hsp70 (Fig. 3A,B; Supplementary Fig. 5A,B).

**Figure 3.**
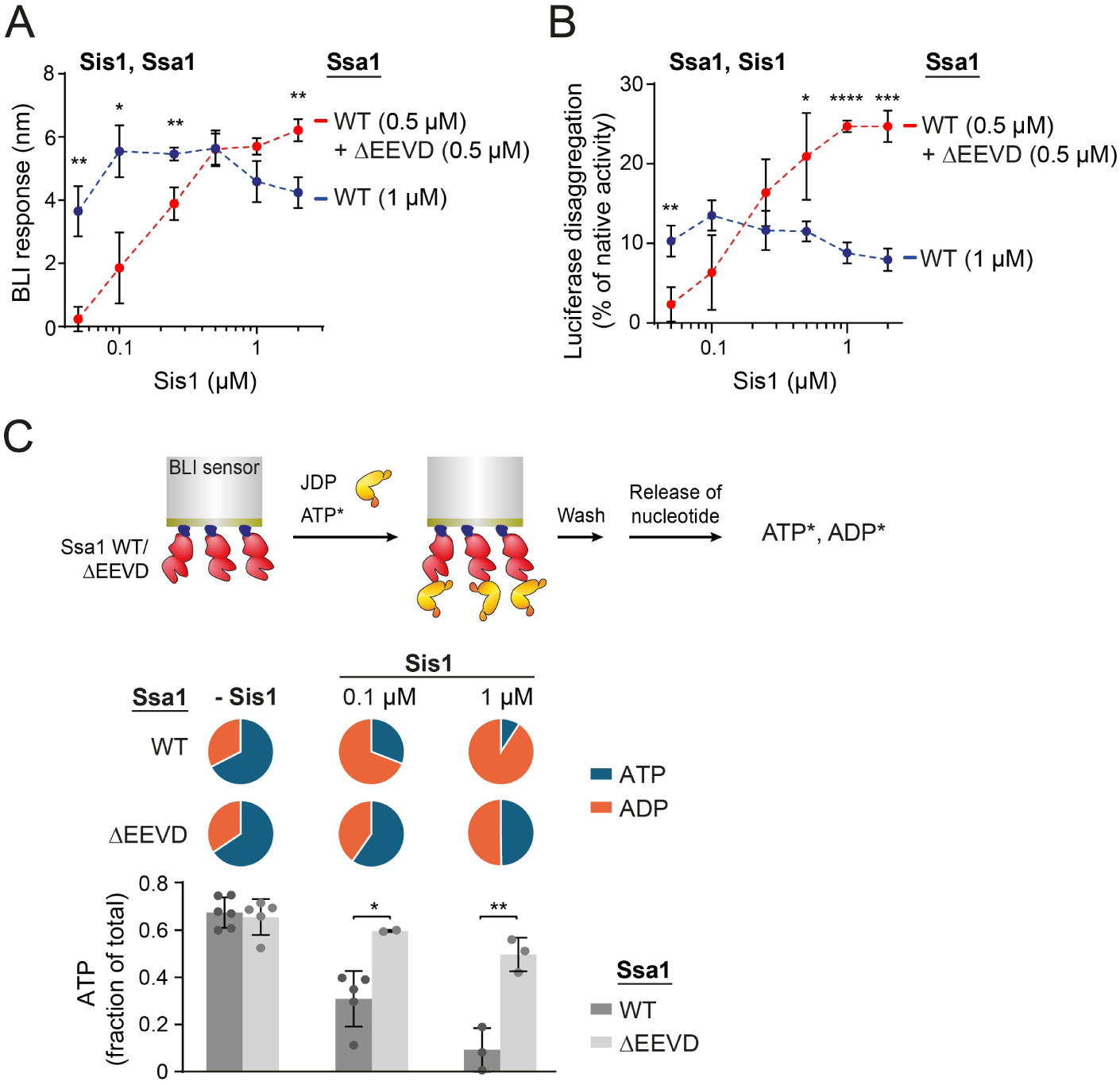
Hsp70-JDP interaction via EEVD sensitizes Hsp70 to inhibition by Class B JDP. (A) Binding of Ssa1^WT^ (blue) or 1:1 mixture of Ssa1^WT^ and Ssa1^ΔEEVD^ (red) to BLI sensor with luciferase aggregates in the presence of the indicated Sis1 concentrations. The plot shows binding signal following incubation with chaperones for 20 minutes. (B) Disaggregation of aggregated luciferase by Ssa1 (blue) or 1:1 mixture of Ssa1^WT^ and Ssa1^ΔEEVD^ (red) in the presence of the indicated Sis1 concentrations. Luciferase activity was measured after 4 h of incubation with chaperones and normalized to the activity of the native protein. (A, B) Shown is mean with SD from at least three repeats. (C) Upper panel: the scheme of the experiment. His-SUMO-tagged Ssa1 variants (Ssa1^WT^ and Ssa1^ΔEEVD^) were immobilized on BLI sensors and incubated with or without Sis1 (concentration indicated in the legend) and [α-P^32^]ATP for 1 h to reach equilibrium. Subsequently, after washing, radioactive nucleotides associated with Ssa1 variants were released and the ratio between ATP and ADP was analyzed by TLC and scintillation counting and presented as pie charts. Bar charts present the same data as ATP fraction of all nucleotides released from Ssa1. Error bars represent SD from the indicated number of repeats. (A-C) Unpaired T test: *p<0.05, **p<0.01, ***p<0.001, ****p<0.0001.

The observed inhibition of Ssa1^WT^ activity by Sis1 might be associated with stabilization of the Ssa1-Sis1 interaction through the EEVD motif. Since the EEVD motif contributes substantially to the formation of the Ssa1-Sis1 complex, deletion of the motif increases the fraction of unbound Ssa1 molecules (Fig. 1 C,D, Supplementary Fig. 1C,D). Accordingly, the observation that Ssa1^WT^ inhibition by Sis1 can be moderated by substituting some Ssa1^WT^ molecules with ΔEEVD variants (Fig. 3A,B) suggests that excessive Hsp70 engagement within the Hsp70-JDP complex may not be optimal for the disaggregation activity. To verify this, we further stabilized the Hsp70-JDP interaction. We took advantage of the fact that the EEVD motif is highly charged and that bivalent ions weaken electrostatic interactions. Consistently, reducing the Mg²⁺ concentration from 15 to 5 mM increased Sis1 binding to Ssa1 (Supplementary Fig. 6A). Interestingly, reducing magnesium ions decreased disaggregation activity of the Ssa1^WT^-Sis1 system and increased the fold stimulation by Ssa1^ΔEEVD^ (Supplementary Fig. 6B). Together, these results demonstrate a correlation between the strength of the Hsp70-JDP interaction, the inhibitory effect of Class B JDPs on Hsp70 activity, and the extent of stimulation by Hsp70^ΔEEVD^.

This raises the question of why the Hsp70-JDP complex stabilized via the EEVD motif, essential for aggregate binding and disaggregation, limits the activity when formed in excess. To explore this, we considered the classical role of JDP in the Hsp70 cycle, in which a JDP first binds a substrate, and together they form a complex with Hsp70, triggering ATP hydrolysis (2). In contrast, Class B JDPs form a complex with Hsp70 through the EEVD motif independently of a substrate and regardless of the nucleotide state of Hsp70 (Fig. 1C,D). This interaction positions the released J-domain in close proximity to Hsp70 and could therefore stimulate ATP hydrolysis in the same Hsp70 molecule that is bound through EEVD. Based on AlphaFold3 models of Ssa1-Sis1 and Hsc70-DNAJB1 complexes, such *in cis* ATPase stimulation is sterically possible (Supplementary Fig. 6C). At high JDP levels, this stimulation might occur in Hsp70 before it engages a substrate, thereby limiting the pool of ATP-bound Hsp70 molecules capable of substrate binding.

To assess whether the EEVD-mediated Class B JDP-Hsp70 interaction may deplete the ATP form of Hsp70, we measured how Sis1 affects the proportion between ATP and ADP bound to Ssa1 and Ssa1^ΔEEVD^. The Ssa1 variants were immobilized on BLI sensors and incubated with or without Sis1 and [α-P^32^]ATP until Sis1 binding reached equilibrium. Subsequently, ^32^P-labeled nucleotides were released from Ssa1 and the ratio between ATP and ADP was assessed (Fig. 3C). In the absence of Sis1, both Ssa1^WT^ and Ssa1^ΔEEVD^ exhibited the same distribution, with approximately 65% of total Ssa1 in the ATP state (Fig. 3C). Upon incubation with Sis1 (1 µM), the ATP-bound Hsp70 fraction decreased to ~10% for Ssa1^WT^ and to ~50% for Ssa1^ΔEEVD^. The impact of Sis1 was concentration-dependent and the presence or absence of the EEVD motif accounted for major differences at different Sis1 concentrations (Fig. 3C).

These results demonstrate that Class B JDP binding to Hsp70 via the EEVD motif decreases the fraction of ATP-bound Hsp70 molecules. Reducing the pool of Hsp70 in a state that can be recruited to aggregates may represent one of the mechanisms underlying the reduced activity of the Hsp70 system at elevated levels of Class B JDPs.

### J-domain autoinhibition moderates negative effects of Hsp70 binding through EEVD

Class B JDP mutants with disrupted J-domain autoinhibition are more potent in stimulating ATP hydrolysis in Hsp70, but do not support Hsp70 clustering on aggregates and disaggregation (22, 23). This resembles the effects observed at elevated JDP concentrations (Fig. 3A,B). Furthermore, as determined by BLI, a Sis1^E50A^ variant with disrupted J-domain autoinhibition (33) showed enhanced binding to sensor-immobilized Ssa1 compared with the wild-type Sis1 (Supplementary Fig. 7A).

To investigate whether the autoinhibition mechanism protects against depletion of ATP-bound Hsp70 upon Class B JDP binding, we quantified the proportions of ATP- and ADP-bound Ssa1 variants with and without the EEVD motif, in the presence of either Sis1^WT^ or Sis1^E50A^. The Sis1^E50A^ variant present at high concentration (1 µM) reduced the proportion of ATP-bound Hsp70 from ~65% to ~5% for both Ssa1^WT^ and Ssa1^ΔEEVD^ (Fig. 4A). However, when the impact was assessed at a lower concentration of Sis1^E50A^ (50 nM), the presence of the EEVD motif was associated with significantly lower proportion of the ATP form: in the case of Ssa1^WT^, it dropped to ~13%, compared to ~50% in the case of the Ssa1^ΔEEVD^ variant. The comparison between Sis^E50A^ and Sis1^WT^ (Fig. 4A) shows that the disruption of autoinhibition substantially reduces the fraction of Hsp70 in the ATP state. Furthermore, the milder effect of Sis^E50A^ on Ssa1^ΔEEVD^ than on Ssa1^WT^ suggests that EEVD binding enhances ATP hydrolysis in Hsp70 by the autoinhibition-defective JDP variant. This is consistent with the hypothesis that ATP hydrolysis can be stimulated *in cis* within the Hsp70-Class B JDP complex formed through the EEVD motif.

**Figure 4.**
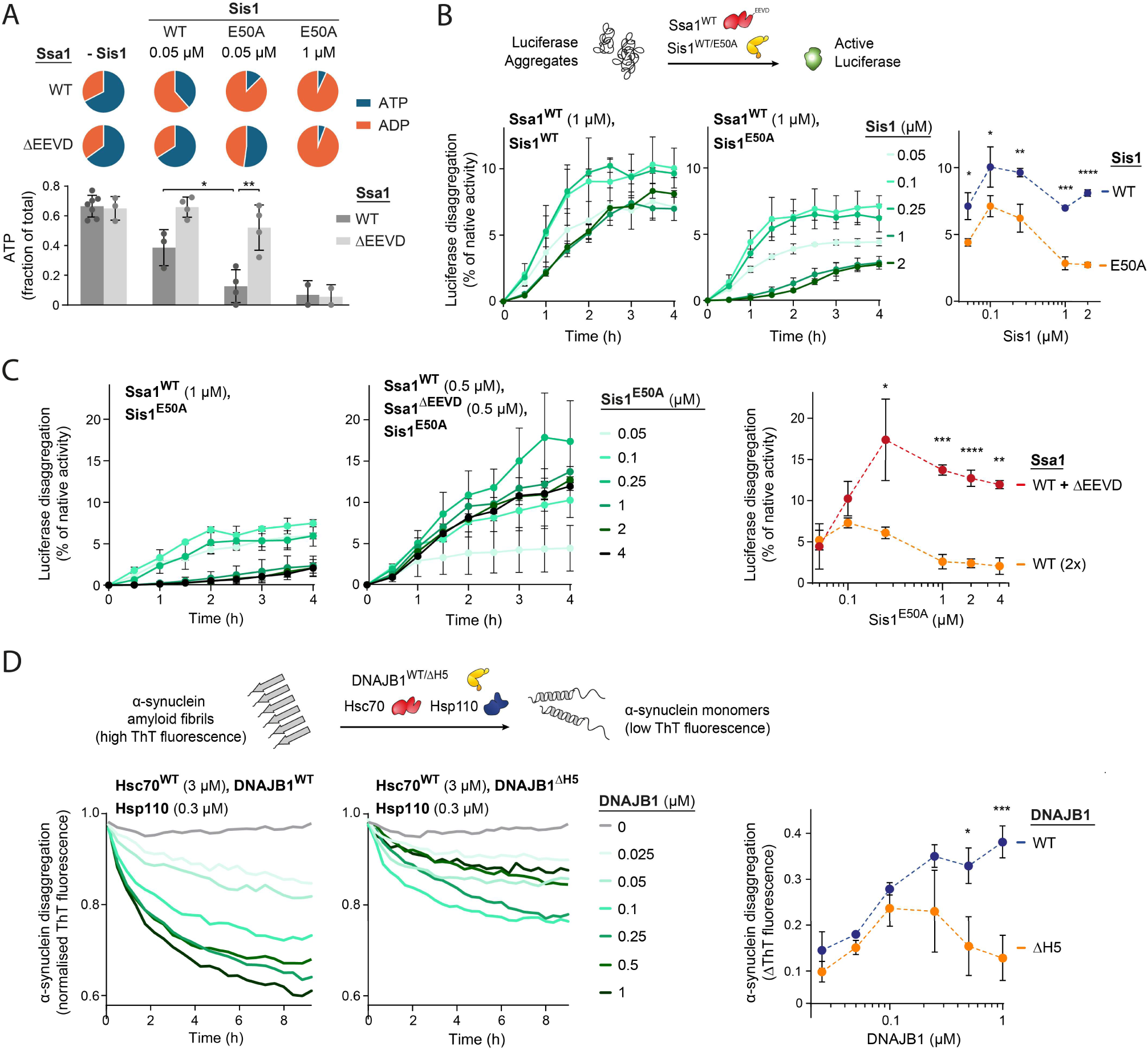
Disruption of J-domain autoinhibition inhibits Hsp70 activity at high JDP concentrations. (A) His-SUMO-tagged Ssa1 variants (Ssa1^WT^ and Ssa1^ΔEEVD^) were immobilized on BLI sensors and incubated with or without Sis1^WT^ or Sis1^E50A^ at indicated concentrations and [α-P^32^]ATP for 1 h to reach equilibrium. Subsequently, the proportion between radioactive ATP and ADP released from Ssa1 was analyzed using TLC and scintillation counting. Results are presented as pie and bar charts. Points indicate the results from independent repeats. Error bars represent SD. Unpaired T test: *p<0.05, **p<0.01. (B) Upper panel: experimental scheme. Disaggregation of aggregated luciferase by Ssa1^WT^ with Sis1^WT^ (left panel) or Sis1^E50A^ (middle panel) present at different concentrations. Activity of luciferase after 4 h of disaggregation (right panel). (C) Disaggregation of aggregated luciferase by Ssa1^WT^ (left panel) or mixture of Ssa1^WT^ and Ssa1^ΔEEVD^ (middle panel) and Sis1^E50A^ present at different concentrations. Activity of luciferase after 4 h of disaggregation (right panel; data from left and middle panels). (D) Upper panel: experimental scheme. Lower panels: disaggregation of preformed α-synuclein fibrils in the presence of Hsc70^WT^ (3 μM), Hsp105 (0.3 μM) and DNAJB1^WT^ or DNAJB1^ΔH5^ at indicated concentrations. Fibrils disaggregation was monitored by Thioflavin T (ThT) fluorescence. Right panel shows change in ThT florescence after 8 h of disaggregation by the indicated chaperones. Fluorescence signal was normalized to the start of the reaction. (B-D) Plots represent mean and error bars represent SD of at least three repeats. Unpaired T test: *p<0.05, **p<0.01, ***p<0.001, ****p<0.0001.

We next asked whether the diminished chaperone activity caused by the disruption of autoinhibition is similar in nature to that observed at high concentrations of wild-type Class B JDP. To investigate this, we tested the ability of the Sis1^E50A^ variant with disrupted autoinhibition to support Ssa1 activity across different JDP concentrations. In both luciferase disaggregation (Fig. 4B) and aggregate-binding assays (Supplementary Fig. 7B), Sis1^E50A^ supported Ssa1 activity only when it was present at low concentrations. At concentrations equimolar to Hsp70 (1 µM) or higher, luciferase disaggregation with Sis1^E50A^ was slower and less productive, and the inhibition was more pronounced than with Sis1^WT^ (Fig. 4B). Interestingly, the addition of the Sse1 nucleotide exchange factor shifted the optimum toward higher Sis1^E50A^ levels (Supplementary Fig. 7C), moderating the inhibitory effect of disrupted autoinhibition.

To determine whether high levels of the autoinhibition-defective JDP variant diminish Hsp70 activity through the interaction with the EEVD motif, we assessed luciferase disaggregation in mixtures of Ssa1^WT^ and Ssa1^ΔEEVD^, in which only half of Hsp70 molecules could engage the JDP via EEVD. Similarly as with Sis1^WT^ (Fig. 3A,B), mixing Ssa1^ΔEEVD^ with Ssa1^WT^ supported disaggregation at high Sis1^E50A^ concentrations much beyond the level achieved with Ssa1^WT^ as the only Hsp70 variant (Fig. 4C). This suggests that the interaction of Class B JDPs with the EEVD motif makes the Hsp70 system substantially more dependent on J-domain autoinhibition.

To test whether the observed trends are conserved, we determined the ability of human DNAJB1^ΔH5^, the variant with distorted Helix 5 responsible for J-domain autoinhibition, to support the disaggregation of α-synuclein fibrils by Hsc70 and Hsp110. Disaggregation was monitored using the Thioflavin T (ThT) assay, in which fibril disassembly results in decreased ThT fluorescence. In contrast to the wild-type DNAJB1, the system comprising DNAJB1^ΔH5^ was active only at low JDP concentrations and was not as effective as the system with DNAJB1^WT^ (Fig. 4D). On one hand, these findings demonstrate that J-domain autoinhibition is not strictly essential for disaggregation of amorphous and fibrillar aggregates. On the other hand, they underscore its protective role against strong inhibition of the Hsp70 chaperone by Class B JDPs, allowing the co-chaperone to support disaggregation across a wider range of concentrations.

Finally, we asked about the role of EEVD binding in the ability of DNAJB1^ΔH5^ to support fibril disassembly. The mixture of Hsc70^WT^ and Hsc70^ΔEEVD^ variants maintained the disaggregation activity at higher DNAJB1^ΔH5^ concentrations than for wild-type Hsc70 variant only and no inhibition was observed up to 2 µM of DNAJB1^ΔH5^ (Supplementary Fig. 8). This further supports the interdependence between EEVD interaction and J-domain inhibition.

In summary, the results suggest that these two Class B-specific traits are functionally coupled. J-domain autoinhibition prevents premature stimulation of ATP hydrolysis – a consequence of Class B JDP interaction with Hsp70 via the EEVD motif. Without this regulatory mechanism, excessive stimulation of ATP hydrolysis by unrestrained J-domains could lead to accumulation of Hsp70 in the ADP-bound state, thereby impairing disaggregation activity of the Hsp70 system.

## Discussion

The finding that the Hsp70 variant lacking the EEVD motif, when added to the wild-type Hsp70 and Class B JDP stimulates the disaggregation (Fig. 2C,D,F) indicates that Hsp70 can be recruited to the aggregate surface via two distinct binding modes: EEVD-dependent and EEVD-independent (Fig. 5). The well-established EEVD-dependent mode is a prerequisite to the EEVD-independent one. It involves JDP binding to Hsp70 through the EEVD motif and release of the J-domain, which activates ATP hydrolysis in Hsp70, resulting in the formation of ultra-affinity complex between the closed SBD of Hsp70 and the aggregate substrate (11, 23, 33, 35). While the J-domain dissociates from Hsp70 in its ADP form, the CTD1 of the JDP retains the interaction with EEVD regardless of the nucleotide- and substrate-binding state of Hsp70 (Fig. 1C,D). This way, due to the EEVD motif, a JDP-Hsp70-aggregate complex can form (Fig. 1E,F). In the second, EEVD-independent, *in trans* binding mode, the activated J-domain, protruding on the extended G/F from the initial JDP-Hsp70-aggregate complex, may serve as a recruiting arm for new Hsp70 molecules to bind to the aggregate in close proximity to the initial complex (Fig. 5).

**Figure 5.**
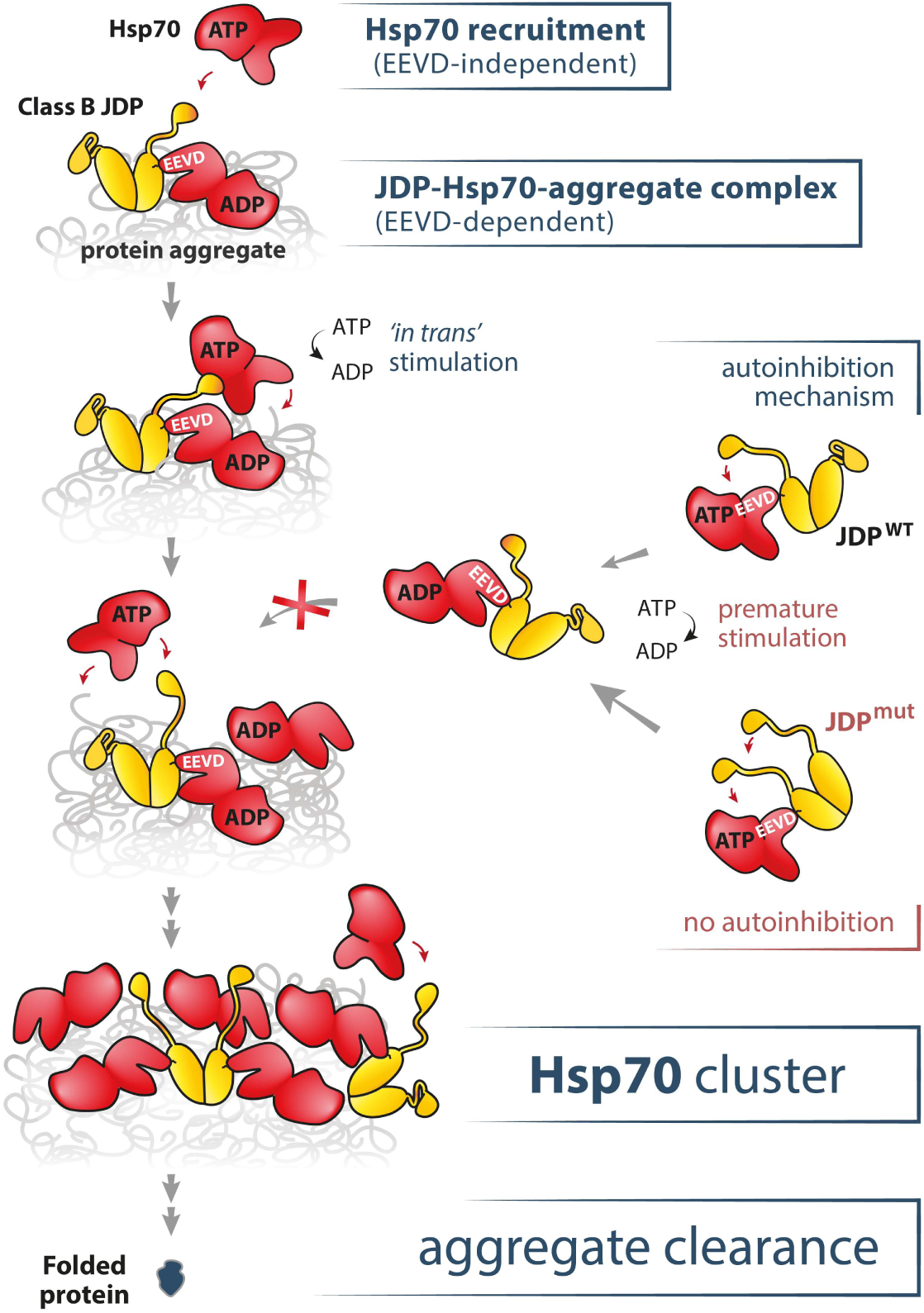
Proposed mechanism of Hsp70 clustering on aggregate surface. In the presence of a Class B JDP, Hsp70 first binds to an aggregate using the EEVD-dependent mode, in which the J-domain is released from autoinhibition due to interaction with the EEVD motif of Hsp70. Subsequently, the activated J-domain of the JDP bound to the Hsp70-aggregate complex may recruit new Hsp70 molecules without engaging their EEVD motifs. The G/F region provides an extended arm for such *in trans* Hsp70 recruitment. Thereby, Class B JDPs support Hsp70 clustering at the aggregate surface, a prerequisite for aggregate dissolution. Right: Thanks to the long G/F, the J-domains can reach the NBD of the same Hsp70 molecule that is bound through EEVD. The stable Class B JDP-Hsp70 interaction via EEVD poses a risk of premature ATP hydrolysis in Hsp70 in the absence of a substrate, yielding Hsp70 in the closed conformation that cannot engage in substrate binding until the rate-limiting nucleotide exchange occurs. The autoinhibition mechanism moderates this negative effect of the EEVD interaction. Lack of autoinhibition mechanism would lead to excessive ATP hydrolysis and depletion of the ATP form of Hsp70, diminishing its disaggregation activity.

Such localized Hsp70 recruitment, constrained by the length of G/F, aligns with the previously described densely packed Hsp70 clusters that form at the tips of alfa-synuclein fibrils (24, 25). These assemblies have been proposed to increase entropic pulling of aggregated polypeptides, thereby facilitating their disaggregation (24, 26), in contrast to the dispersed, non-effective binding supported by Class A JDPs (23). Accordingly, the herein proposed multi-step Hsp70 clustering mechanism sheds light on how the traits specific to cytosolic/nuclear Class B JDPs, namely binding to the Hsp70 EEVD motif and the longest G/F region amongst all Class A and B JDPs (16), contribute to their exceptional ability to support Hsp70 clustering and effective protein disaggregation.

However, the inhibition observed at increased JDP concentrations shows that these properties of Class B JDPs come at a price; since the complex between JDP and Hsp70 via EEVD is stable even in the absence of a protein substrate (Supplementary Figs. 1B,C,D, 7A), leaving the J-domain activated, futile ATP hydrolysis may occur in Hsp70 before it binds to an aggregate (Fig. 5). This is supported by the observation that the Sis1-Ssa1 interaction via EEVD increases the proportion of the ADP state of Ssa1 (Fig. 3C). This effect is exacerbated upon the disruption of J-domain autoinhibition in the Sis1 E50A variant (Fig. 4A), which raises the possibility that the autoinhibition mechanism had evolved to compensate for the negative effects of the stable Class B JDP-Hsp70 interaction through the EEVD motif. Furthermore, the finding that J-domain autoinhibition is required specifically in the context of EEVD binding clarifies why Class A JDP, lacking both these features, does not inhibit Hsp70 in Class B JDP-driven protein disaggregation, but the two co-chaperones cooperate, making Hsp70 a more potent disaggregase (47, 48). An important question for future studies is how the formation of mixed Class A/B JDP complexes (47, 48), as well as the presence of other chaperones on aggregate surface further influence Hsp70 functional cycle.

The observation that wild-type Ssa1 harboring the EEVD motif is much more sensitive to the mutation compromising J-domain autoinhibition than Ssa1^ΔEEVD^ (Fig. 4A) also suggests that ATP hydrolysis can be stimulated within the same Hsp70 molecule that is bound through EEVD (*in cis*) (Fig. 5; Supplementary Fig. 6C). This provides an additional mechanism of ATPase stimulation alongside the well-stablished one, whereby a substrate-bound JDP triggers ATP hydrolysis in an Hsp70 molecule engaged with the same substrate – a mechanism proposed also for Class A JDPs and Class B DNAJB6, which do not interact with EEVD (23, 37, 49). In contrast, *in cis* ATPase stimulation may occur both on aggregate surface (Fig. 2G) and prematurely, in the absence of a substrate (Fig. 5). Such premature ATP hydrolysis, combined with slow nucleotide exchange (13) may decrease what can be defined as the effective Hsp70 concentration – comprising molecules in an ATP-bound, open conformation capable of substrate binding. Without the autoinhibition mechanism, unrestrained J-domains locally concentrated within the Hsp70-JDP complex stabilized via EEVD may trigger premature ATP hydrolysis already at low JDP concentrations. Consequently, autoinhibition-deficient JDP mutants could reduce the effective Hsp70 concentration below the threshold required for formation of disaggregation-competent Hsp70 clusters on aggregates (Fig. 5).

Given this, a question arises why the addition of the Hsp70 ^ΔEEVD^ variant, which maintains a substantially higher proportion of the ATP-bound state than the wild-type (Fig. 4A), stimulates aggregate binding and disaggregation (Fig. 2A,C,D,F), yet it does not promote folding of a non-aggregated substrate (Fig. 2E)? Similarly, why mutations that disrupt J-domain autoinhibition specifically impair aggregate disassembly but not protein folding (22, 23)? Extending the hypothesis formulated by Wentink and co-workers (2020), we speculate that Hsp70 clustering on aggregates may require higher effective Hsp70 concentration than the canonical dispersed binding, or binding to a non-aggregated misfolded protein. One possible explanation is the limited number of chaperone-binding sites available in the vicinity of the initial Hsp70–JDP-aggregate complex, together with the relatively weak affinity of Hsp70 for the J-domain/G-F recruiting arm. Further studies will be needed to test whether Hsp70-driven disaggregation is indeed more susceptible to premature ATP hydrolysis than protein refolding. Overall, our results raise the possibility that the dynamics and topology of Hsp70 cycle may differ between protein disaggregation and refolding activities.

The negative impact on the effective Hsp70 concentration may not be the only factor limiting the disaggregation activity upon disruption of J-domain autoinhibition. Problems may also arise due to other postulated roles of this regulatory mechanism, such as maintaining correct Hsp70 distribution on the aggregate (49) and driving coordinated amyloid disassembly (25). Furthermore, ordered amyloids and amorphous aggregates may have different requirements for these functions. Nonetheless, it is curious that the effects of mutations such as E50A in Sis1 or disruption of the inhibitory Helix 5 in DNAJB1 can be partially rescued simply by reducing the level of the JDP variant (Fig. 4B,D). This suggests that the problems caused by the lack of autoinhibition may be similar in nature to those observed at higher concentrations of the wild-type JDP. Thus, it is feasible that the autoinhibition mechanism allows Class B JDPs to be produced in the cell at sufficiently high levels to support their other physiological functions, such as folding or preventing aggregation of a specific substrate (e.g. PIKKs and tau, physiological substrates of Sis1(50) and DNAJB1(51)) while still supporting Hsp70-mediated disaggregation.

The impairment of disaggregation caused by the lack of autoinhibition can also be overcome by Hsp110 (Supplementary Fig. 7C), as also demonstrated by Faust et al. (2020) for the human Hsc-DNAJB1-Hsp110 in α-synuclein disassembly. Previous studies suggested that the primary role of the NEF in disaggregation is its ability to selectively recycle Hsp70, thanks to its substrate-release function biased toward more accessible non-clustered Hsp70 molecules (24, 25, 52). Our results suggest that also the nucleotide exchange function itself may improve disaggregation by Hsp70, otherwise inhibited e. g. due to excessive *in cis* ATP hydrolysis associated with the stable Class B JDP-Hsp70 interaction via EEVD (Fig. 5). All these mechanisms are not mutually exclusive, and the clustering-driven disaggregation may be regulated by the co-chaperones at multiple levels.

At the same time, it should be noted that NEFs strongly destabilize Hsp70-substrate interactions (2, 3, 53). Although elevated NEF concentrations may restore the disaggregation activity of Hsp70 in the presence of autoinhibition-deficient Class B JDP mutants, the same concentrations inhibit disaggregation supported by the wild-type JDP (Supplementary Fig. 7C). Therefore, increasing Hsp110 expression as a strategy to compensate for defects associated with J-domain autoinhibition should be considered with caution.

Overall, our findings indicate that the evolutionary innovations in JDP co-chaperones that potentiated the Hsp70 disaggregase against amyloids and amorphous aggregates are more interdependent than previously assumed. Understanding this regulatory framework is especially important in the light of the ongoing attempts to selectively modulate the interaction network between Hsp70 and JDPs as a therapeutic approach against neurodegenerative diseases and chaperonopathies involving aberrant DNAJB4, DNAJB6 and DNAJB2 variants (36, 37, 54). Due to the bi-phasic impact of Class B JDP on Hsp70, boosting the disaggregation of amyloids such as α-synuclein, Tau or amyloid-β, may require either strengthening or disrupting the Hsp70-JDP interactions, depending on the physiological levels of Hsp70 and co-chaperones. In line with that, recent promising use of a small-molecule inhibitor of the Hsp70 JDP interaction against mutations that disrupt autoinhibition in DNAJB6 associated with LGMDD1 myopathy (37), triggers an urgent need to understand mechanistical similarities and differences between EEVD-interacting (canonical, e.g. DNAJB1) and non-interacting Class B paralogs (non-canonical, e.g. DNAJB6). The gain-of-function character of myopathy-linked mutations in Class B JDPs highlights the ability of these co-chaperones to compromise broad Hsp70 functionality (36, 37). A potential mechanism may involve decreasing the level of Hsp70 in the ATP-bound, open state (Fig. 4A). In the future, understanding molecular basis of this vulnerability may guide the development of therapeutic strategies that restore proteostasis in disorders associated with Hsp70 and JDP dysfunction.

## Materials and Methods

### Proteins

Published protocols were used for purification of Ssa1 (14), Ssa1^ΔEEVD^ (14), Sis1 (22), Ydj1 (55), α-synuclein (16) and the His-SUMO construct (22). Hsc70, Hsc70^ΔEEVD^, DNAJB1, DNAJB1^ΔH5^, and Hsp105 were purified following procedures described by (47). His-tagged firefly luciferase was obtained as previously reported (56), whereas untagged luciferase was purchased from Promega (E1701). All Ssa1 variants (Ssa1^T201A^, Ssa1^V435F^, Ssa1^ΔEEVD^) as well as Sis1^E50A^ and Sis1^1–338^ were generated by PCR-based site-directed mutagenesis (Qiagen) and verified by DNA sequencing. The His-SUMO-Ssa1 variant was expressed and purified analogously to Ssa1, except that the proteolytic removal of the His-SUMO tag by Ulp1 protease was omitted to retain the fusion tag. Protein concentrations were quantified for monomers, using densitometry with Albumin Standard (Thermo Scientific). Protein concentrations are given for monomers.

### Protein labeling

Ssa1 was labeled with the Alexa Fluor 594 C5-Maleimide Labeling Kit (ThermoFischer Scientific) and Ssa1 T201A and Ssa T201A ΔEEVD and Ssa ΔEEVD were labeled with the Alexa Fluor 488 C5-Maleimide Labeling Kit (ThermoFischer Scientific) according to the manufacturer instructions, using a protein:dye ratio 1:20, in magnesium acetate buffer (50 mM HEPES pH 7.5, 100 mM KCl, 5 mM Mg(CH_3_COO)_2_, 10% Glycerol) with 1 mM TCEP. After labeling, extensive dialysis was performed to remove the unbound dye.

### Bio-Layer Interferometry (BLI)

Bio-layer interferometry measurements were performed using BLItz or Octet K2 instruments (Sartorius) using Octet NTA Biosensors (Sartorius) at 25 °C with 2000 rpm mixing according to previously published protocols (22, 52, 56). Statistical analyses were performed using GraphPad Prism 6 software.

#### BLI: Hsp70-JDP binding

*Sensor preparation:* His-SUMO-Ssa1 WT or His-SUMO-Ssa1^ΔEEVD^ (2 µM) in the HKM buffer, pH 7.5 (25 mM HEPES-KOH, 75 mM KCl and 15 mM MgCl_2_, if not stated otherwise) was loaded onto the NTA BLI sensor for 1 min, reaching 2.5 nm of the BLI signal, followed by washing with the buffer supplemented with ATP (5 mM, if not stated otherwise) for 30 mins. Subsequently, sensors were incubated with Sis1 variants used at concentrations indicated in figure legends and washed with the same buffer lacking Sis1 to monitor dissociation (Supplementary Fig. 1A).

For experiments with the ϕ peptide (5’TAMRA-FYQLALT, GenScript), presented in Fig. 1C, D and Supplementary Fig. 1B, the NTA BLI sensor with immobilized Hsp70 or Hsp70^ΔEEVD^ was incubated in the HKMGTDA buffer pH 7.5 (25 mM HEPES-KOH, 75 mM KCl, 15 mM MgCl_2_, 5% glycerol, 0,002% Tween 20, 1 mM DTT, 1 mM ADP or ATP), supplemented with (or without) the ϕ peptide (10 µM) for 30 minutes to allow Hsp70 loading with the peptide. Tween 20 was included to prevent nonspecific binding of ϕ peptide to the sensor. Subsequently, the sensor was incubated in the HKMGTDA buffer containing JDP (0.4 µM for Sis1 and 2 µM for DNAJB1, or no JDP,) in the presence or absence of the ϕ peptide (10 µM) (JDP association step) and then transferred into the buffer with or without the ϕ peptide (JDP dissociation step).

To control for Hsp70 binding to the substrate in the presence of ADP, free His-SUMO-Hsp70 (1 µM) was incubated with the ϕ peptide (10 µM) for 30 minutes under the same conditions as in the BLI assay. Subsequently, the mixture (75 µl) was loaded onto ZEBA Spin Desalting Columns 7K MWCO (Thermo Scientific) and the complex was separated from the unbound peptide via size exclusion chromatography (void volume elution 66 µl), according to the manufacturer’s protocol. Concentration of the co-eluted ϕ peptide divided by concentration of eluted Hsp70 constituted 98.3%, 79.5% and 99.6% (for His-SUMO-Ssa1; results of 3 independent repeats) and 81.7%, 57% and 93,1% (for His-SUMO-Hsc70; results of 3 independent repeats). The concentration of eluted Ssa1 was calculated using densitometry, and the concentration of the co-eluted peptide was calculated based on fluorescence intensity measurement, with subtracted background fluorescence of the void volume elution fraction in the absence of Hsp70. Fluorescence was measured using Tecan Spark 10M Multimode Microplate Reader (excitation 545 nm, emission 590 nm).

#### BLI: Binding to luciferase aggregates

*Sensor preparation:* NTA BLI biosensor was equilibrated with the HKM buffer pH 8.0 (25 mM HEPES-KOH, 75 mM KCl, 15 mM MgCl_2_) for 10 min and subsequently incubated in the buffer supplemented with 6 M urea and His-tagged luciferase (8.2 µM) to allow formation of the initial anchoring protein layer of ~6 nm (Supplementary Fig. 2). Subsequently, the sensor was immersed in a solution containing native luciferase (1.6 µM) in HKM buffer and incubated at 44 °C for 10 minutes. During this step, luciferase misfolds and binds to urea-unfolded luciferase on the sensor resulting in the formation of luciferase aggregates with a thickness of approximately 15– 20 nm. The sensor was subsequently equilibrated with the HKM buffer supplemented with 5 mM ATP and 2 mM DTT.

Binding and dissociation of chaperones were monitored in the buffer HKM, pH 8.0, with 5 mM ATP and 2 mM DTT. Chaperones were used at concentrations indicated in figure legends.

For the BLI experiments presented in Fig. 1E, F, the baseline, chaperone binding, and dissociation steps were performed in binding buffer pH 7.5 (25 mM HEPES-KOH, 75 mM KCl, 5 mM MgCl2, 2 mM DTT, 5 mM ATP and 0.002% Tween-20). Tween 20 was included to prevent protein binding the sensor outside the bio-layer. The yeast system binding was performed for 1200 s., followed by dissociation step for 600 s. The human system binding was performed for 5400 s., followed by dissociation step for 1800 s. Length of the binding step was determined by saturation of the binding signal. Ssa1, Ssa1 DEEVD and Sis1 were used at 1 μM concentration. Hsc70, Hsc70 ΔEEVD and DNAJB1 were used at 2 μM concentration.

### Western Blot

For the analysis of aggregate-bound chaperones on the BLI sensor, the baseline and chaperone binding, were performed in the binding buffer (25 mM HEPES-KOH pH 7.5, 75 mM KCl, 5 mM MgCl_2_, 2 mM DTT, 5 mM ATP and 0.002% Tween-20). Hsc70, Hsc70^ΔEEVD^ and DNAJB1 were used at 2 μM concentration. The binding step was performed for 5400 s, without the dissociation

Sensor stripping was performed upon the end of the BLI run by submerging the sensor tip in the Laemmli sample buffer (Thermo Fisher) with 50 mM EDTA and incubating at 95 °C for 5 minutes. Next, the samples were loaded on 12% SDS-PAGE gel and the electrophoresis was performed in a running buffer containing 25 mM Tris-HCl, 192 mM Glycine and 1% SDS, under fixed voltage (150 V) for 1.5 hours. Proteins were transferred from gels onto nitrocellulose blotting membranes (Cytiva). The membranes were subsequently blocked with 3% milk in TBS-T (20 mM Tris, 150 mM NaCl, 0.1% Tween) for 30 minutes, washed with TBS-T and incubated with primary antibodies against the indicated proteins for 1 hour at room temperature, followed by incubation with appropriate HRP-conjugated secondary antibodies (1 h, room temperature). All blots were analyzed with a Chemidoc Imager (BioRad) and quantified in ImageJ (NIH). Statistical analyses were performed using GraphPad Prism 6 software.

The following antibodies were used: Hsp70 (5A5, ab2787, abcam, 1:1000 dilution), DNAJB1 (13174-1-AP, Proteintech, 1:10 000 dilution), Ssa1 (A302-519A, Thermofischer Scientific, 1:600 dilution), Sis1 (COP-COP-080051, Cosmo Bio Co., 1:5000 dilution), goat anti rabbit IgG-HRP (BioRad), goat anti mouse IgG-HRP (BioRad).

### Luciferase disaggregation

Disaggregation of firefly luciferase was performed according to two established protocols previously used for the characterization of yeast (22) and human (12) chaperone systems, which differed in aggregate preparation, chaperone concentrations, and reaction buffer composition. In the setup using yeast chaperones, luciferase at a final concentration of 30 µM was chemically denatured in HKM buffer pH 8.0 (25 mM HEPES–KOH, 75 mM KCl, 15 mM MgCl₂) supplemented with 8 M urea. The protein was incubated for 10 min at 25 °C and subsequently transferred to 48 °C for an additional 10 min. Denatured luciferase was then rapidly diluted 25-fold into the same buffer to induce aggregation. Following the 25-fold dilution, luciferase aggregates were incubated for 4 min at room temperature before being added to the reaction mixtures at a final concentration of 50 nM together with the indicated chaperones. All reactions were carried out in HKM buffer, pH 8.0, supplemented with 5 mM ATP and 2 mM DTT at 25 °C, except for experiments assessing the effect of MgCl₂ concentration, where ATP concentration was 1 mM. In the setup using human chaperones, luciferase (2 µM) was chemically denatured in buffer comprising 40 mM HEPES–KOH pH 7.5, 50 mM KCl, 5 mM MgCl₂ supplemented with 6 M urea. The protein was incubated at 30 °C for 30 min, following 2-fold dilution with buffer lacking chaperones to induce aggregation. Aggregates were subsequently diluted 50-fold into disaggregation buffer (40 mM HEPES–KOH pH 7.5, 50 mM KCl, 5 mM MgCl₂, 2 mM DTT, 5 mM ATP, 3 mM PEP, 20 ng/µl pyruvate kinase, 10 µM BSA) containing the indicated chaperone proteins and incubated at 30 °C. Luciferase reactivation was quantified using the Luciferase Assay Kit (Promega, E1501) and luminescence was measured using a GloMax instrument (Promega). Statistical analyses were performed using GraphPad Prism 6.

### Unfolded luciferase reactivation

To assess the spontaneous folding of luciferase, the method was adapted from (57). Luciferase (10 μM) was denatured by incubation in 5 M guanidine hydrochloride (GuHCl) supplemented with 10 mM dithiothreitol (DTT) at 25 °C for 1 h. The refolding reaction was initiated by the rapid 100-fold dilution into the buffer comprising 25 mM HEPES-KOH pH 7.5, 100 mM KCl, 10 mM Mg(OAc)_2_, 5 mM ATP, 2 mM DTT, 0.05% Tween 20, and chaperones indicated in figure legends. Luciferase reactivation was quantified using the Luciferase Assay Kit (Promega, E1501) and luminescence was measured using a GloMax instrument (Promega).

### α-synuclein disaggregation

α-synuclein fibrils were prepared as described (16, 24). Briefly, monomeric α-synuclein (200 μM) was incubated for 15 days in 20 mM HEPES-KOH pH 7.5, 100 mM NaCl at 37 °C, with shaking (1,000 rpm). Fibrils were concentrated by centrifugation (20,000×g, 30 min) and resuspended in 20 mM HEPES-KOH pH 7.4, sonicated and stored at −80 °C. Fibril formation was confirmed using Thioflavin T (ThT) fluorescence, measured in the in a 96-well half-area nonbinding microplate (Corning 3881) using Tecan Spark plate-reader (excitation: 440 nm, emission: 480 nm).

Disaggregation of α-synuclein was monitored as described (16), with small modifications. Preformed α-synuclein fibrils (0.8 μM, monomer) were incubated at 30 °C in 25 mM HEPES-KOH pH 7.5, 75 mM KCl, 15 mM MgCl2, 2 mM DTT, 2 mM ATP, with ATP regeneration system (8 mM phosphoenolpyruvate and 86 nM pyruvate kinase, Sigma) and 30 μM ThT. After the addition of chaperones: 3 μM Hsc70, 0.3 μM Hsp105 and DNAJB1 or DNAJB1^ΔH5^ at the concentrations 25, 50, 100, 250, 500 or 1000 nM, the reaction mixtures were incubated for 15 min to reach 30 °C and the fluorescence was measured for 10 h. The signal for proteins without fibrils was subtracted.

### ATP/ADP quantification

Ni-NTA BLI sensor (Sartorius) was incubated with 2 μM His-SUMO-Ssa1 or His-SUMO-Ssa1ΔEEVD in the HKM buffer (25 mM HEPES-KOH pH 7.5, 75 mM KCl, 15 mM MgCl_2_), for 15 min, washed, and incubated with or without Sis1 or Sis1^E50A^ (concentrations indicated in the figures) in the HKM buffer supplemented with 0.1 mM ATP with 50 μCi/ml [α-P^32^]ATP (Hartmann Analytic) for 1 h at room temperature. An advantage of such approach over the classical chromatography-based methods of separation of Hsp70 in complex with nucleotides is the ability to release nucleotides from Hsp70 within few seconds after the incubation with Sis1 and [α-P^32^]ATP, minimizing ATP hydrolysis that would otherwise occur before enzyme inactivation. After 4 instant washes by submerging the Sis1-immobilised sensor in the HKM buffer, the nucleotides were released into 2 M acetic acid and separated on the Silica TLC plates coated with PEI-modified cellulose with Fluorescent Indicator F254 (Sigma) with 1 M formic acid/1 M LiCl. ATP:ADP ratio was quantified using Beckman LS 6000 Liquid Scintillation Counter. To correct for the background nucleotide binding to the sensor, nucleotide level bound to sensor-immobilized SUMO-His protein was subtracted. Statistical analyses were performed using GraphPad Prism 6 software.

## ACKNOWLEDGMENTS

This study was supported by National Science Centre, Poland grant (OPUS 27 2024/53/B/NZ1/00632). The work of D.P., K.K. and B.T. was supported by a National Science Centre, Poland grant (OPUS 21 2021/41/B/NZ8/02835). We thank Dr. Anne Wentink and Dr. Rina Rosenzweig for plasmids for JDP and α-synuclein expression.

