## Supplementary Information for "Aggregate-Bound Chaperone Complexes Promote Disaggregation through J-Domain-Driven Hsp70 Recruitment"

**This PDF file includes:**

Supporting text
Supplementary Figures 1 to 8
SI References

### Materials and Methods

**Bio-Layer Interferometry:** ***Chaperone Binding to aggregated yeast lysate.*** *Sensor preparation:* Yeast lysate was prepared from an overnight culture of W303 cells grown in YPD medium, following the procedure described by (Wyszkowski *et al*, 2021). Ni-NTA biosensors were first hydrated with the buffer HKM, pH 8, for 10 min and subsequently incubated for 10 min in the buffer HKM, pH 8, containing 6 M urea and 8.2 µM His-tagged luciferase to form an anchoring layer. After a 5-min wash with the buffer, the biosensors were transferred into the buffer supplemented with 5 mg/mL soluble yeast proteins and incubated at 55 °C for 15 min. The sensors were then equilibrated with the buffer containing 5 mM ATP and 2 mM DTT until the biolayer thickness reached approximately 30 nm. Binding and dissociation of chaperones were monitored at 25 °C in the buffer HKM, pH 8.0, with 5 mM ATP and 2 mM DTT. Chaperones were used at concentrations indicated in figure legends.

**Microscale Thermophoresis**

Binding experiments were performed with the Monolith NT.115 instrument (NanoTemper Technologies). For binding experiments, 200 nM of Alexa Fluor 488-labeled Ssa1 variants (Ssa1 T201A and Ssa T201A ΔEEVD) were mixed (1:1 volumes, final concentration 100 nM) with a range of Sis1 serial dilutions (starting from 26 µM) in the MST buffer pH 8 (50 mM HEPES-KOH, 50 mM KCl and 0,002% Tween-20), and incubated at room temperature for 10 min. Samples were centrifuged at 15 000 RCF for 10 min before loading into standard-treated MST capillaries (NanoTemper Technologies). The measurements were performed at room temperature with 40% excitation power and 60% MST power. All measurements were performed at least three times. The collected MST data were analyzed using MO Affinity Analysis Software (NanoTemper Technologies).

**AlfaFold predictions of Hsp70-JDP complexes**

For Hsp70-JDP complexes modeled with the AlphaFold 3 server (Abramson *et al*, 2024), we used full-length Hsp70, in combination with full-length JDP and the CTD2 domain, to model dimer orientation. For the Ssa1-Sis1 complex (seed: 735456653, ipTM = 0.62, pTM = 0.57), input sequences included: wild-type Ssa1 (Uniprot ID P10591), wild-type Sis1 and CTD2 of Sis1 (265-351) (Uniprot ID P25294). For the Hsc70-DNAJB1 complex (seed: 1974609510, ipTM = 0.57, pTM = 0.57), input sequences comprised: wild-type Hsc70 (Uniprot ID P11142), wild-type DNAJB1 and CTD2 of DNAJB1 (251-340) (Uniprot ID P25685).

### Supplementary Figures

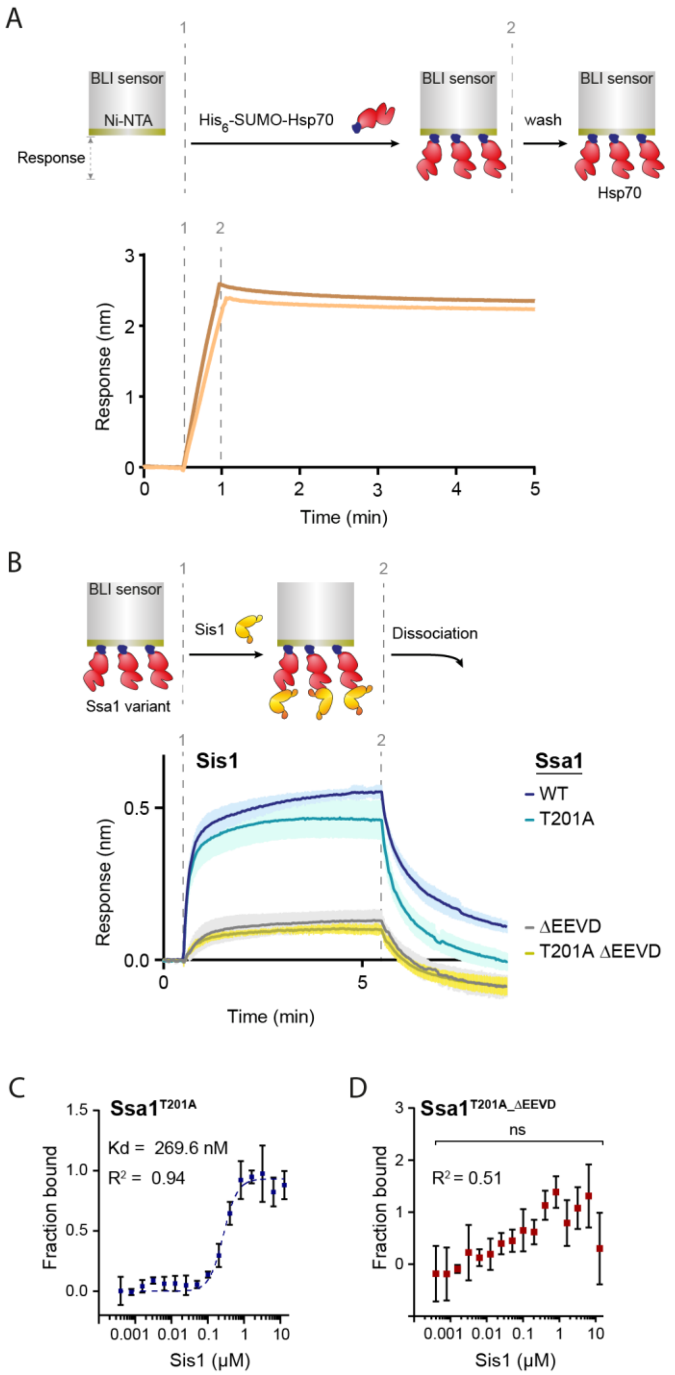


**Supplementary Fig. 1.** (A) Preparation of BLI sensors with immobilized Hsp70 variants. Upper panel shows the sensor preparation workflow. Dashed line 1 indicates start of $His_6$-SUMO-Hsp70 variant binding to a Ni-NTA BLI sensor, dashed line 2 indicates wash of the sensor. Lower panel shows BLI signal from preparation of two sensors with $His_6$-SUMO-Ssa1. Hsp70 loading was carried out until the signal reached approximately 2.5 nm to yield similar amount of immobilized protein on each sensor. (B) Upper panel: experimental scheme. Interaction of Ssa1 and its variants with Sis1 in the presence of ATP. Ssa1 variant (WT, T201A, ΔEEVD or T201A ΔEEVD) was immobilized on the BLI sensor as presented in panel A, following incubation with Sis1 (0.4 µM). Binding curves represent the mean of three replicates, with shaded areas indicating SD. Dashed line 1 indicates the start of the Sis1 binding step, dashed line 2 indicates the dissociation step into buffer lacking chaperones. (C, D) Affinity between Ssa1$^{T201A}$ and Sis1 (C), Ssa1$^{T201A_\Delta EEVD}$ and Sis1 (D), assessed by microscale thermophoresis. Ssa1$^{T201A}$ and Ssa1 $^{T201A\_\Delta EEVD}$ were labeled with Alexa Fluor 488. The Ssa1$^{T201A}$ variant was used to maintain Ssa1 in the ATP-bound state. Error bars represent SD from three repeats.

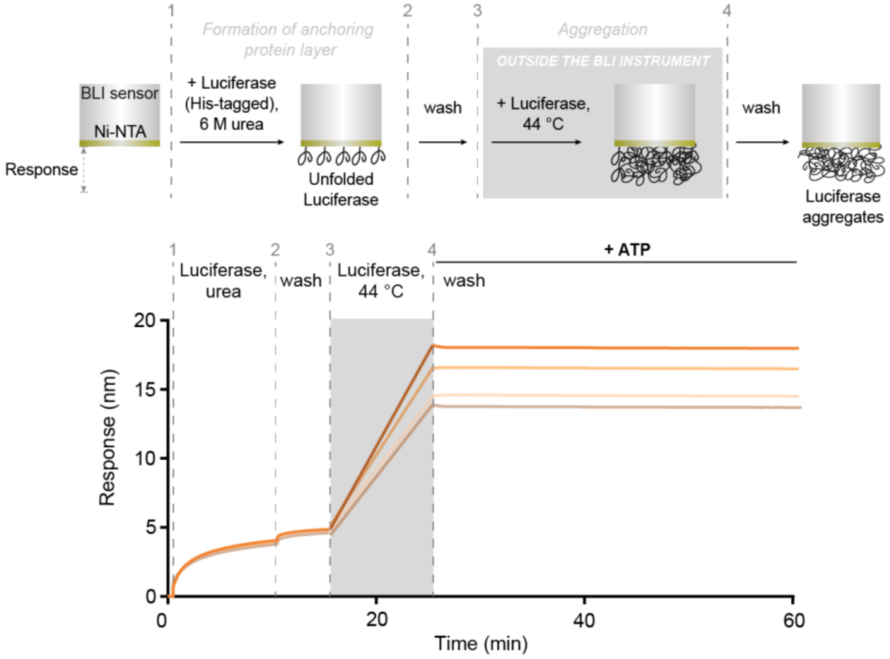


**Supplementary Fig. 2.** Preparation of BLI sensors with luciferase aggregates. Dashed lines indicate the steps of the workflow. Ni-NTA BLI sensors were first incubated with urea-unfolded His-tagged luciferase at 25 °C to form an anchoring protein layer (1), following wash with buffer (2) and subsequent incubation with luciferase at 44 °C to allow aggregate formation (3). BLI response was not recorded during the step 3. Thickness and stability of aggregate layer was monitored with BLI during final washing (4). BLI traces represent four independent preparations of BLI sensors with luciferase aggregates.

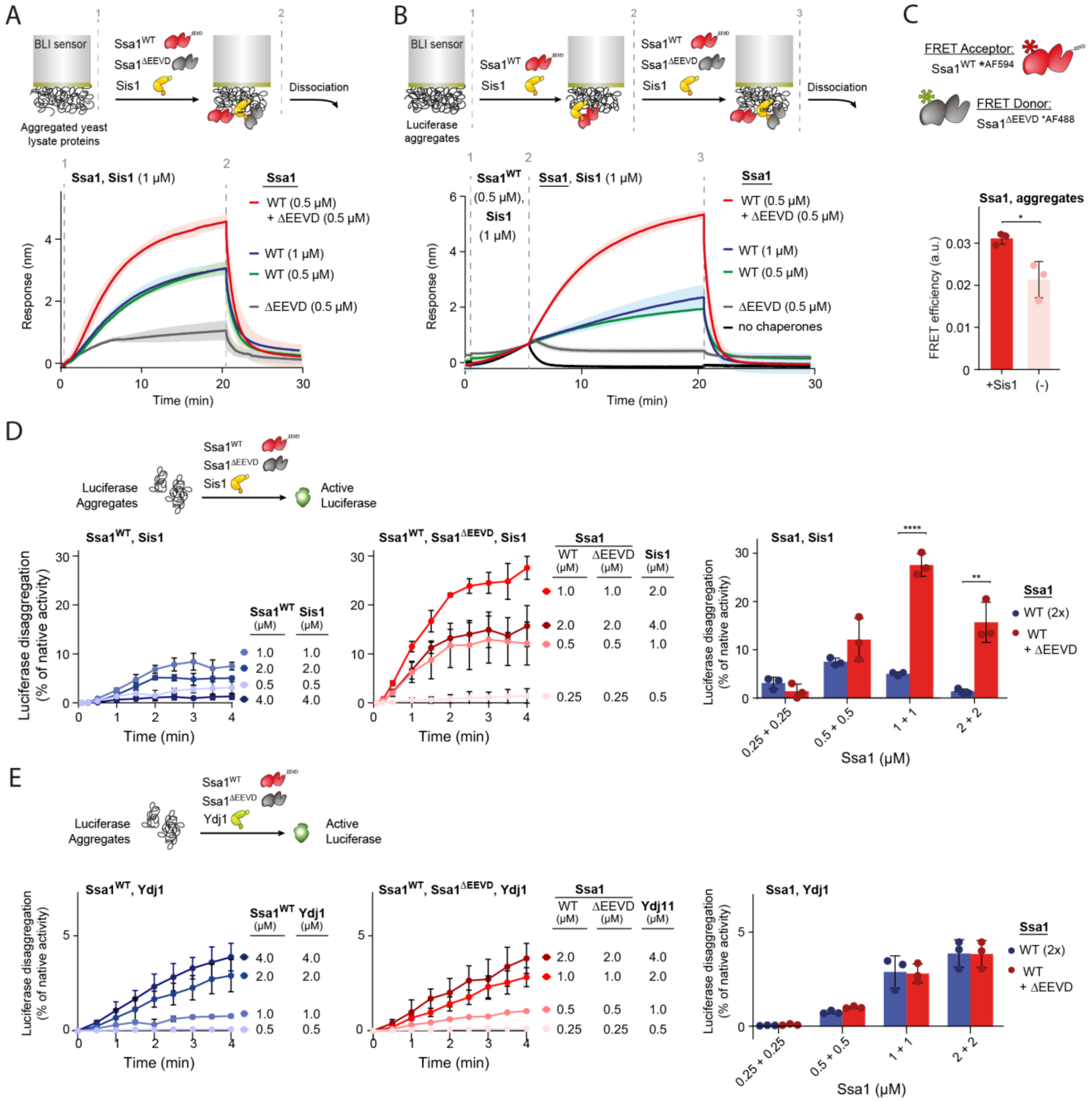


**Supplementary Fig. 3.** (A) Ssa1 variant lacking the EEVD motif (Ssa1$^{\Delta EEVD}$) stimulates chaperone binding to aggregates when Ssa1$^{WT}$ and Sis1 are present. Upper panel: experimental scheme. BLI sensor with immobilized aggregated proteins from yeast lysate was incubated with Sis1 and Ssa1$^{WT}$ or Ssa1$^{\Delta EEVD}$ or 1:1 mixture of WT Ssa1 and Ssa1$^{\Delta EEVD}$, as indicated in the legend. Dashed line 1 indicates the start of the binding step, dashed line 2 indicates the dissociation step into buffer lacking chaperones. (B) Ssa1$^{\Delta EEVD}$ binding to aggregates requires the presence of the Ssa1$^{WT}$ variant. Upper panel: scheme of sequential binding of the indicated chaperones to sensor-immobilized luciferase aggregates. Dashed line 1 indicates the start of the initial incubation with the Ssa1$^{WT}$-Sis1 system, common for all traces, dashed line 2 indicates the start of subsequent incubation with Sis1 and Ssa1 variants indicated in the legend. Dashed line 3 indicates the start of the dissociation step. (A, B) Binding curves represent the average of three repeats, shades designate SD. Control without chaperones during the second incubation (black) is shown as a signal from two repeats. (C) Quantification of FRET efficiency between Ssa1$^{WT}$ AF594 (acceptor) and Ssa1$^{\Delta EEVD}$

AF488 (donor) (0.5 µM each) and luciferase aggregates (0.25 µM) in the presence and absence of Sis1 (0.5 µM). Unpaired T-test: * $p<0.05$. (D) Upper panel: experimental scheme. Lower panels: disaggregation of aggregated luciferase by Sis1 (concentration equal to that of Ssa1) and $Ssa1^{WT}$ or a 1:1 mixture of $Ssa1^{WT}$ and $Ssa1^{\Delta EEVD}$ as indicated in the legend. Right panel: luciferase activity following 4 hours of disaggregation. Unpaired T test **$p<0.01$, ****$p<0.0001$. (E) In the presence of Class A JDP Ydj1, luciferase disaggregation with $Ssa1^{\Delta EEVD}$ is not stimulated above the level of $Ssa1^{WT}$. Upper panel: experimental scheme. Lower panels: disaggregation of aggregated luciferase by Ydj1 (concentration equal to that of Ssa1) and $Ssa1^{WT}$ or a 1:1 mixture of $Ssa1^{WT}$ and $Ssa1^{\Delta EEVD}$ as indicated in the legend. Right panel: luciferase activity measured following 4 hours of disaggregation. (D, E) Luciferase activity was measured at indicated timepoints and normalized to the activity of native luciferase. (C, D, E) Data represent the mean ± SD of three independent repeats.

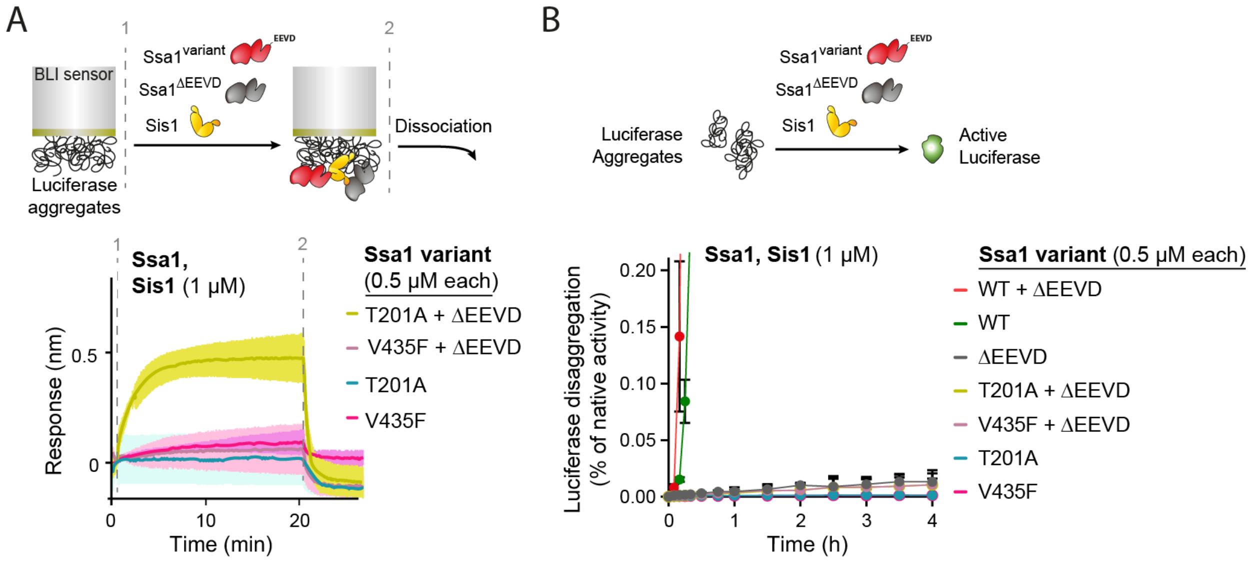


**Supplementary Fig. 4**. Functional requirements for EEVD-independent Hsp70 recruitment to aggregates. (A, B) Upper panel: experimental scheme. (A) Role of ATP hydrolysis and substrate binding in Ssa1 in the recruitment of Ssa1$^{\Delta EEVD}$ to aggregates. BLI sensor with aggregated luciferase was incubated with Sis1 and the indicated Ssa1 variants. Curves represent the average of three repeats; shades indicate SD. Dashed lines indicate chaperone binding (1) and dissociations (2) steps. (B) Disaggregation of aggregated luciferase by Sis1 and the indicated variants Ssa1. Luciferase activity was measured at the indicated time points and normalized to the activity of native luciferase. Error bars represent SD from three repeats.

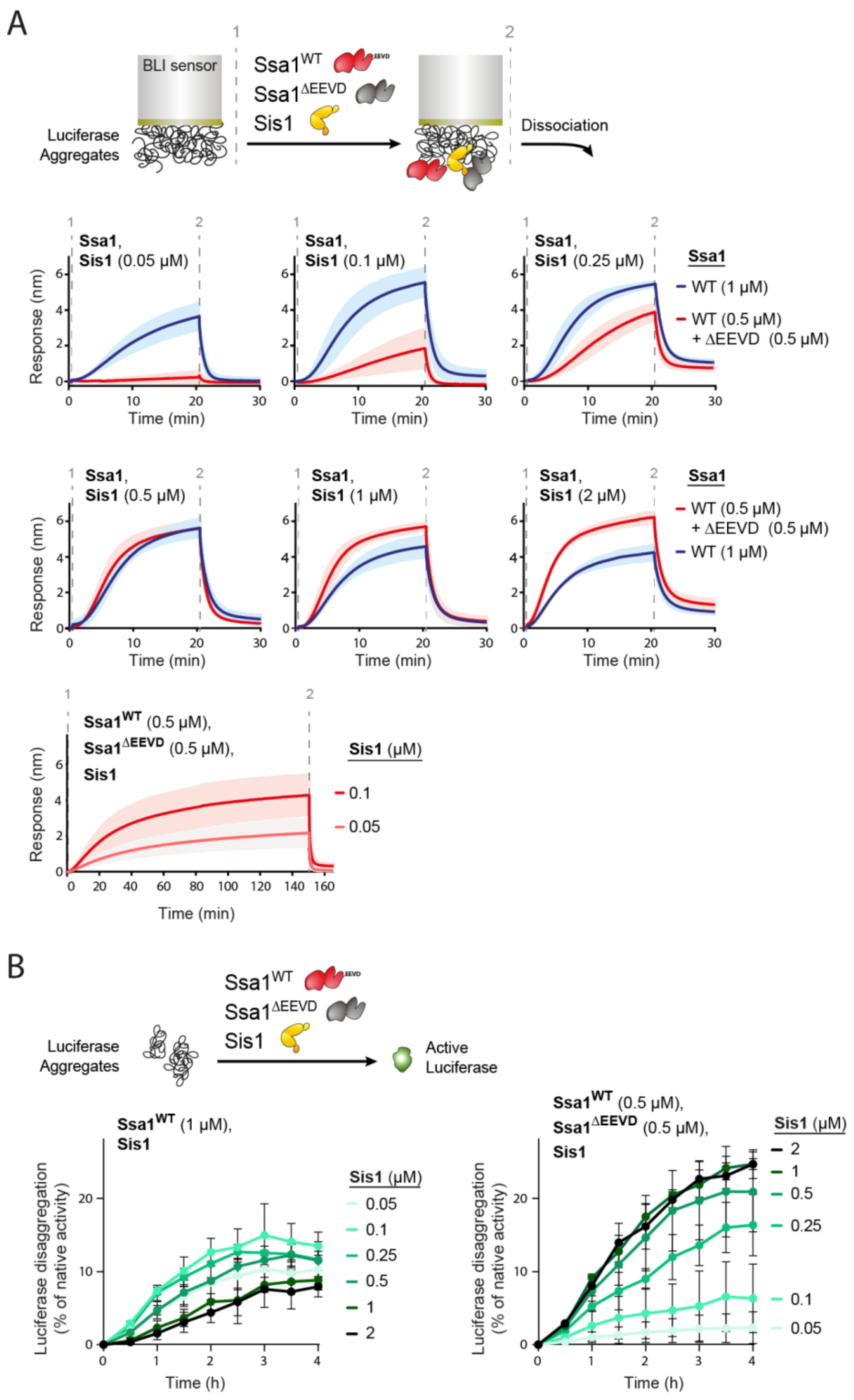


**Supplementary Fig. 5.** (A) Binding of Ssa1$^{WT}$ (blue) or 1:1 mixture of Ssa1$^{WT}$ and Ssa1$^{\Delta EEVD}$ (red) to BLI sensor with luciferase aggregates in the presence of the indicated Sis1 concentrations. Dashed line 1 indicates the start of the binding step, dashed line 2 indicates the dissociation step into buffer lacking chaperones. The bottom panel shows results with extended binding time for the mixture of Ssa1$^{WT}$ and Ssa1$^{\Delta EEVD}$ at the lowest Sis1 concentrations. Binding curves represent the average of three repeats, shades designate SD. (B) Disaggregation of aggregated luciferase by Ssa1 (left) or 1:1 mixture of WT Ssa1 and

$Ssa1^{\Delta EEVD}$ (right) in the presence of the indicated Sis1 concentrations. Luciferase activity was measured at the indicated time points and normalized to the activity of native luciferase. Error bars represent SD from three repeats. (A, B) Upper panel, experimental workflow.

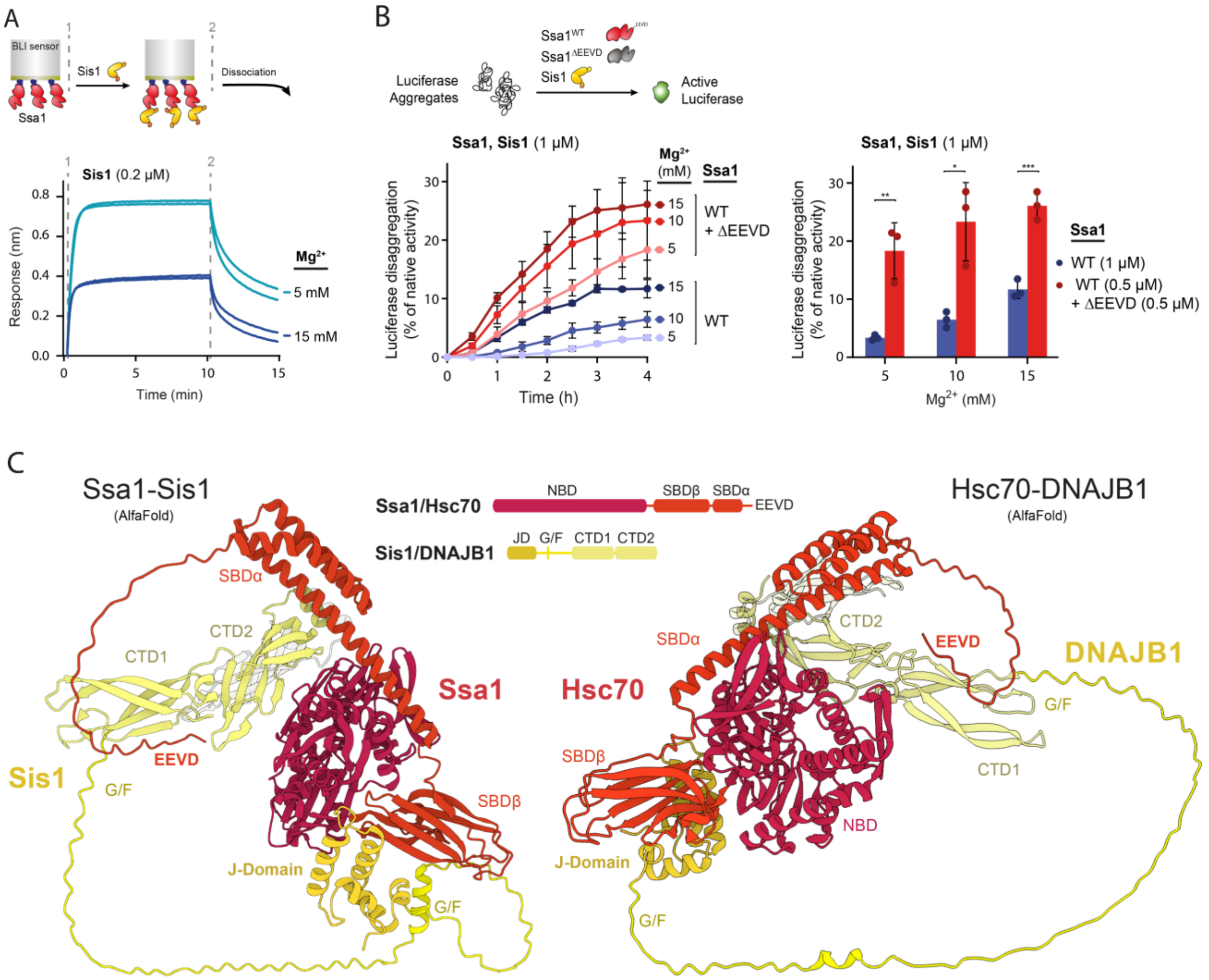


**Supplementary Fig. 6.** (A) BLI sensor with immobilized $His_6$-SUMO-Ssa1 was incubated with Sis1 in the presence of 5 mM or 15 mM magnesium chloride and 1 mM ATP. BLI traces show signal from 2 repeats. (B) Upper panel, scheme of the experiment. Disaggregation of aggregated luciferase by Sis1 and Ssa1 variants: $Ssa1^{WT}$ or $Ssa1^{\Delta EEVD}$, or 1:1 mixture of WT Ssa1 and $Ssa1^{\Delta EEVD}$, under the indicated concentrations of magnesium ions. Right panel: luciferase activity following 4 hours of disaggregation. Luciferase activity was measured at indicated timepoints and normalized to the activity of native luciferase. Error bars represent SD from three repeats. (A, B) Upper panel: scheme of the experiment. (C) Left panel: AlfaFold3 model of full-length Ssa1 in complex with Sis1 dimer comprising one full-length Sis1 and the CTD2 domain of the second Sis1 monomer (residues 265-351) (grey). Right panel: AlfaFold3 model of full-length Hsc70 and DNAJB1 dimer comprising one full-length DNAJB1 monomer and the CTD2 domain of the second DNAJB1 monomer (251-340) (grey). CTD2 domains were introduced to account for dimerization in JDP orientation relative to Hsp70. Middle panel: color coded schematic presentation of Ssa1/Hsc70 and Sis1/DNAJB1 domains.

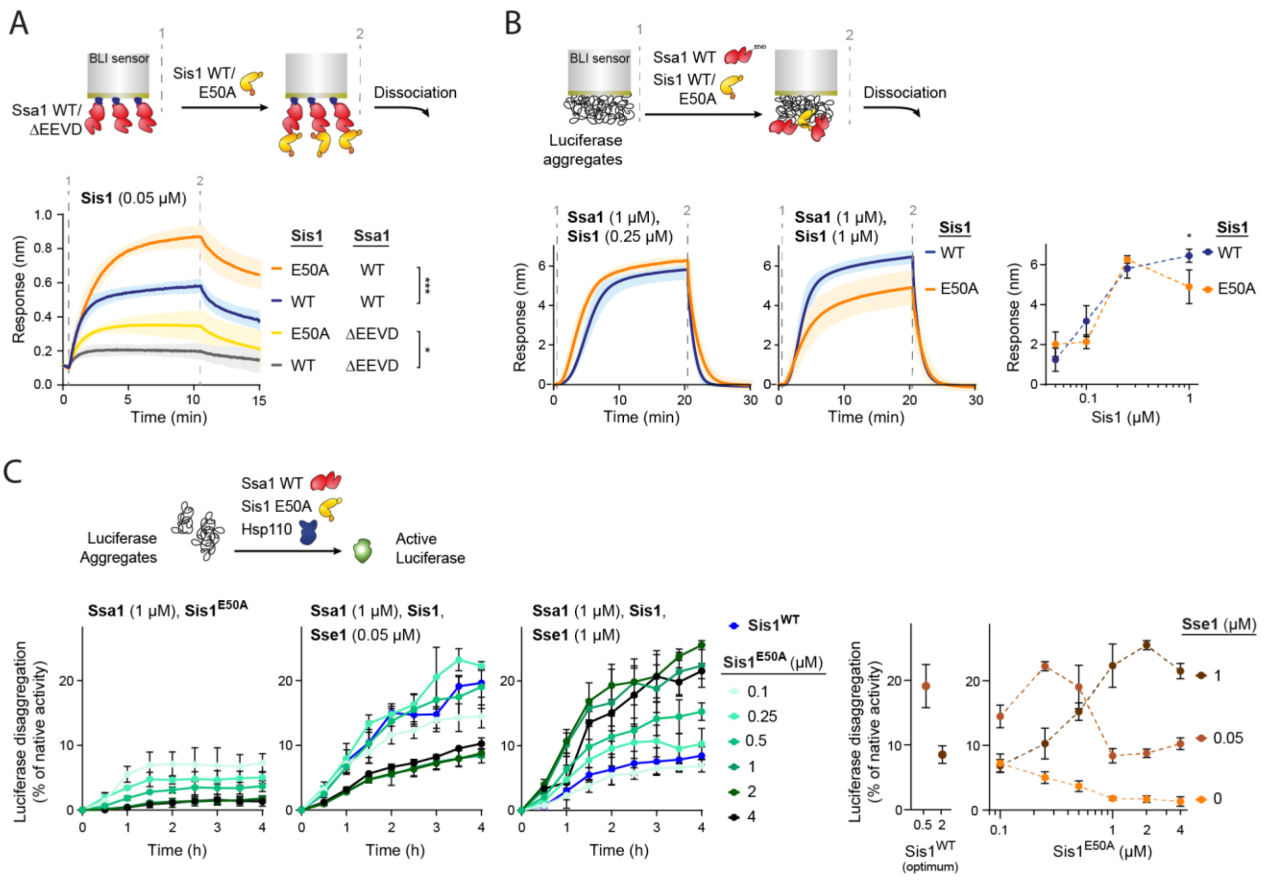


**Supplementary Fig. 7.** (A) BLI sensor with immobilized Ssa1$^{WT}$ or Ssa1$^{\Delta EEVD}$ was incubated with Sis1$^{WT}$ or Sis1$^{E50A}$. (B) BLI sensor with aggregated luciferase was incubated with Ssa1 and Sis1 variants: Sis1$^{WT}$ or Sis1$^{E50A}$ at the indicated concentrations. Right panel: signal following 20 minutes of incubation with the chaperones was plotted as a function of Sis1 concentrations. Data for 0.25 µM and 1 µM Sis1 are from the left and middle panels. (A, B) Binding curves represent the average of three replicates, shades and error bars designate SD. Unpaired T test: *$p<0.05$, ***$p<0.001$ (C) Disaggregation of aggregated luciferase by Ssa1 and Sis1$^{E50A}$, without Sse1 (left panel) or with Sse1 (middle panels). As a reference, disaggregation with Ssa1 and Sis1$^{WT}$ is shown in blue, with Sis1$^{WT}$ used at concentration yielding maximal disaggregation activity (optimum) at the indicated Sse1 concentrations (0.5 µM Sis1$^{WT}$ at 0.05 µM Sse1 and 2 µM Sis1$^{WT}$ at 1 µM Sse1). Luciferase activity was measured at the indicated time points and normalized to the activity of the native protein. Right two panels: normalized luciferase activity after 4 h of disaggregation corresponding to the datasets presented in the three panels on the left. Error bars represent SD from three repeats. (A-C) Upper panels: scheme of the experiment.

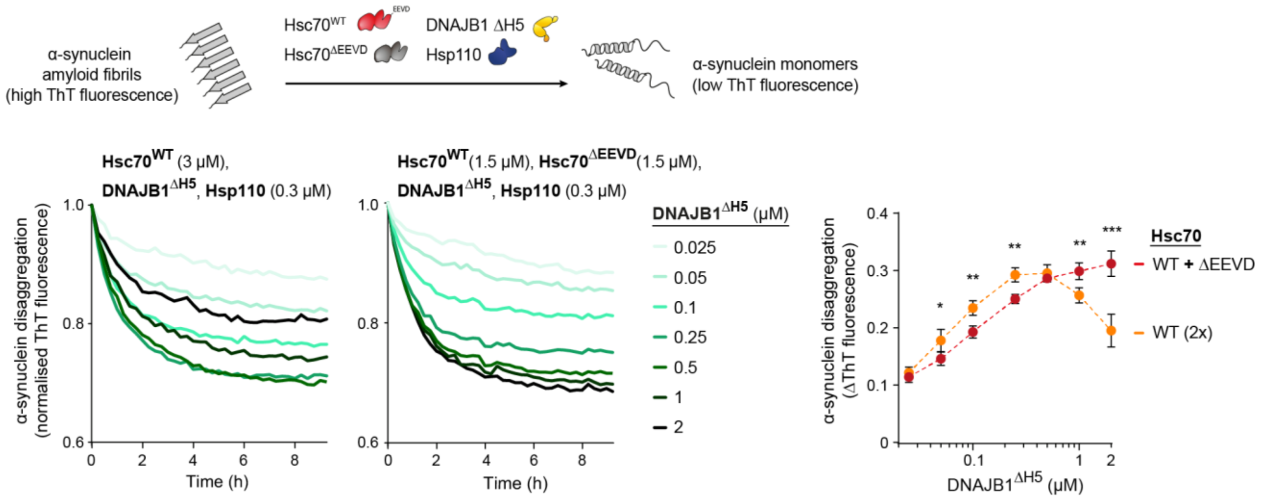


**Supplementary Fig. 8.** Disaggregation of preformed α-synuclein fibrils, in the presence of $Hsc70^{WT}$ (left panel) or mixture of $Hsc70^{WT}$ and $Hsc70^{\Delta EEVD}$ (middle panel) and Hsp105 (0.3 μM) and $DNAJB1^{\Delta H5}$ at indicated concentrations. Right panel shows change in ThT florescence after 8 h of disaggregation by chaperones. Fluorescence signal was normalized to the start of the reaction. Plots represent mean and error bars represent SD of at least three repeats. Upper panel: scheme of the experiment. Unpaired T test: *p<0.05, **p<0.01, ***p<0.001.
